# Apical size and *deltaA* expression predict adult neural stem cell decisions along lineage progression

**DOI:** 10.1101/2022.12.26.521937

**Authors:** Laure Mancini, Boris Guirao, Sara Ortica, Miriam Labusch, Felix Cheysson, Valentin Bonnet, Minh Son Phan, Sébastien Herbert, Pierre Mahou, Emilie Menant, Sébastien Bedu, Jean-Yves Tinevez, Charles Baroud, Emmanuel Beaurepaire, Yohanns Bellaiche, Laure Bally-Cuif, Nicolas Dray

**Author notes:** Laboratoire Reproduction et Développement des Plantes, ENS de Lyon, Lyon, France. LAMA, Université Gustave Eiffel, UMR CNRS 8050, Champs-sur-Marne, France. Imaging Core Facility, Biozentrum, University of Basel, Basel, Switzerland (Sébastien Herbert).

## Abstract

The maintenance of neural stem cells (NSCs) in the adult brain depends on their activation frequency and division mode. We use long-term intravital imaging of NSCs in the zebrafish adult telencephalon to link activation and division mode with predictive cellular and molecular parameters. We reveal that apical surface area and expression of the Notch ligand DeltaA predict NSC activation frequency, while *deltaA* expression marks NSC commitment to neurogenesis. We also find that *deltaA*-negative NSCs constitute the *bona fide* self-renewing NSC pool and systematically engage in asymmetric divisions generating a self-renewing *deltaA*^*neg*^ and a neurogenic *deltaA*^*pos*^ NSC. Finally, modulation of Notch signaling during imaging indicates that the prediction of activation frequency by apical size, and the asymmetric divisions of *deltaA*^*neg*^ NSCs, are functionally independent of Notch. These results provide dynamic qualitative and quantitative readouts of NSC lineage progression *in vivo* and support a hierarchical organization of NSCs in differently fated sub-populations.

## Introduction

Neural Stem Cells (NSCs) produce neurons and glial cells important for the physiology and plasticity of the adult vertebrate brain^1–3^. To ensure these functions, NSC populations remain active and neurogenic during adult life. The efficiency of NSC population maintenance varies greatly between species and with age, and the mechanisms involved remain incompletely understood.

NSC population maintenance is the net result of two major fate decisions: NSC activation from quiescence (over time leading to NSC exhaustion), and the occurrence of self-renewing divisions. These decisions are actively studied in the telencephalic niches of the mouse and zebrafish adult brain (sub-ependymal zone -SEZ-of the lateral ventricle and sub-granular zone -SGZ-of the hippocampus in mouse, pallium in zebrafish), today the most tractable and comparable models to address NSC behavior *in vivo*^4,5^. In both species, NSCs reside most of their time in the G0 quiescence state, under control of quiescence-promoting pathways (notably Notch2/3 and BMP signaling) that gate the frequency of G1 entry (so-called “activation”) *in vivo*^6^. These factors are superimposed to cell-intrinsic windows of responsiveness to activation signals^7^, which remain to be identified. Concerning division mode, clonal tracing and intravital imaging in mouse and zebrafish highlight that NSCs can divide in a symmetric self-renewing (NSC/NSC), asymmetric (NSC/NP) or symmetric neurogenic (NP/NP) fashion where NP (“TAP” -transit amplifying progenitor-in mouse) is a non-glial intermediate progenitor committed to neuron generation after a few divisions. The balance between these distinct outcomes affect NSC maintenance amplification, steady-state and loss, respectively^8–15^. These choices could be cell-autonomous or involve some degree of cell-cell interactions, which could explain the coordination observed at the level of the population^14,16,17^. *In vitro*, the NSC/NP division mode of adult mouse NSC involves the asymmetric expression of the Dyrk1a kinase, a regulator of Notch signaling^18,19^. When the Notch ligand Delta-like1 is overexpressed, asymmetric NSC/NP division also correlates with the segregation of Delta-like1 in the NP daughter^20^. *In vivo*, NSC populations at any time form a patchwork of asynchronous NSCs with distinct molecular states^21–24^, morphologies^25,26^ and fate^10,14,27,28^ and regulators of NSC fate decisions within the intact neurogenic niche remain to be identified.

To identify mechanisms controlling NSC activation and fate decisions *in vivo*, we used intravital imaging to reconstruct adult NSC lineages and decipher cell-intrinsic features that characterize NSC decisions. Adult NSCs in the zebrafish adult pallium are radial glial cells, intermingled with NPs, covering the pallial surface. Thanks to this superficial position, one can record the behavior of all progenitor cells (NSCs and NPs) in their intact neurogenic niche, with single cell resolution and over weeks^8,15,29,30^. Thus, lineages can be captured at long-term, allowing to monitor their progression in spite of very slow time dynamics^8,30^. To characterize pallial NSC lineage features, we performed long-term (up to 52 days) intravital imaging of the NSC population in a double transgenic context expressing a transcriptional reporter of *deltaA*^31^, the most expressed Notch ligand in the adult pallium^32^, and a novel reporter line for tight junctions to highlight apical NSC shapes. We optimized previous methods and analyzed about 1000 NSC tracks in situ, under physiological conditions or upon manipulation of apical surface area (AA) and Notch signaling, eventually allowing to link AA and *deltaA* expression with self-renewal, neurogenic potential and lineage progression.

## Results

### Cellular hallmarks quantitatively correlating with NSC states

Cell geometry regulates essential processes such as growth, lineage commitment, and signaling in various stem cell systems^33,34^. Additionally, apical cell shape in embryonic epithelia correlate with or determine cell fate^35–37^. To characterize the organization of NSCs within their niche, we first assessed which cell state/type possesses an apical contact. We focused on the dorsal (Dm) and anterior (Da) pallial areas, which have been most extensively analyzed and are actively engaged in neurogenesis^8^. We performed triple whole-mount immunohistochemistry (IHC) on adult pallia (3 months post-fertilization -mpf-) from the *Tg(gfap:eGFP)* transgenic line^38^ to label the tight junction protein Zona Occludens 1 (ZO1) together with NSCs (GFP+) and cell proliferation (Proliferating Cell Nuclear Antigen, PCNA) (Figure 1A). This combination identifies quiescent NSCs (qNSCs) (GFP+, PCNA-), activated NSCs (aNSCs) (GFP+, PCNA+), and activated NPs (aNPs) (GFP-, PCNA+), while ZO1 delimits the apicobasal interface of polarized cells. Of note, PCNA is visible throughout the G1-S-G2-M phases of the cell cycle and shortly after the division event, resulting in doublets of aNSCs^8,29^; thus, we only selected isolated aNSCs (aNSC singlets) to focus on aNSCs prior to division. We found that all three cell types display apicobasal polarity and contact the pallial ventricle through their apical surface. They are also in direct contact with each other through their ZO1-positive tight junction, forming a monolayer. Besides, we qualitatively observed that the nuclei of qNSCs and aNSCs are very close to and aligned with the pallial ventricular surface, whereas aNP nuclei are often found in a deeper position (Figure 1B), in line with aNPs being on their way to delaminate upon differentiation into neurons. These observations highlight that the adult pallial germinal pool is organized as a pseudo-stratified neuroepithelial monolayer where the apical surfaces of qNSCs, aNSCs and aNPs are spatially intermingled.

**Figure 1.**
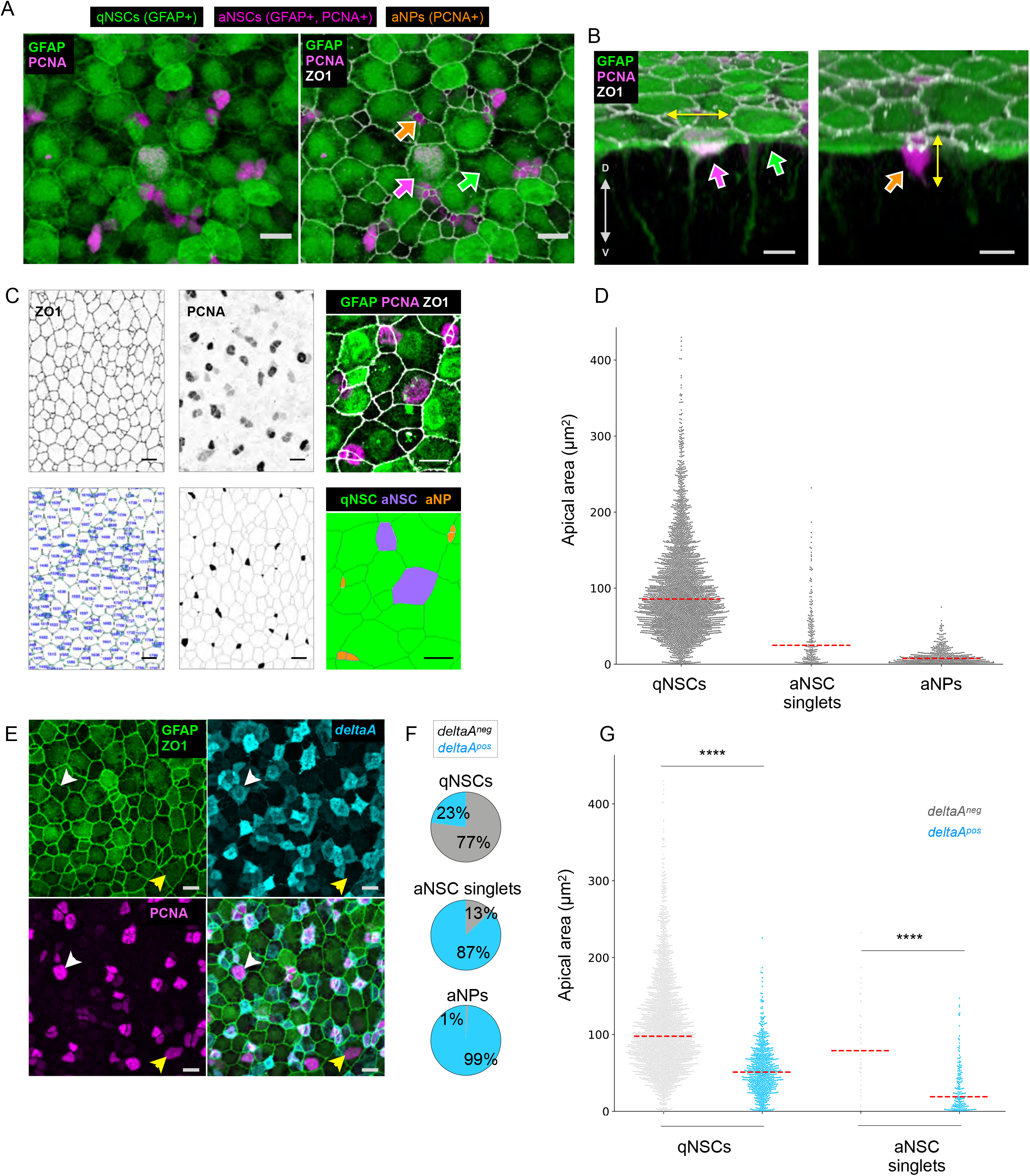
Apical area correlates with cellular types and states in the germinal zone of the adult zebrafish pallium. **1A-1B**. High magnification of a 3-mpf *Tg(gfap:eGFP)* whole-mount pallium processed for immunohistochemistry (IHC) for GFP (green), PCNA (magenta) and ZO1 (white). Arrowheads point towards qNSCs (GFP+, green), aNSCs (GFP+ PCNA+, magenta) and aNPs (PCNA+, orange). **A**. Dorsal view. ZO1 reveals the apical area (AA) of each cell and the cellular topology of the tissue. **B**. Optical section perpendicular to the pallial ventricular zone, passing, from left to right, through a qNSC, an aNSCs and a cluster of two aNPs. Double-headed white arrow to the direction of the dorsoventral axis, double headed yellow arrows to the main axis of elongation of the nucleus in an aNSC and an aNP. Scale bars: 10μm. **1C**. Top: photographs of whole-mount pallia (dorsal views, Dm); bottom: corresponding segmented views (green circle: cell vertex). Left: segmentation of AA based on ZO1 (blue number: Cell ID used for subsequent analysis). Middle: Segmented regions with detected marker, here PCNA (manually corrected segmentation to avoid false-positive cells). Right: High magnification of a preparation stained for the markers of interest with the corresponding final segmentation and cell identities (color coded). **1D**. Distribution of AAs of qNSCs, aNSCs and aNPs in Dm for 4 brains pooled. Red dashed lines: median. **1E**. Maximum projection of a dorsal view of a whole-mount IHC in a 3-mpf *Tg(deltaA:eGFP);Tg(gfap:dtTOMATO)* fish labelled for dTomato (green), ZO1 (green), eGFP (cyan) and PCNA (magenta). Arrowheads to a *deltaA*^*pos*^ aNSC (white) and *deltaA*^*neg*^ aNSC (yellow). Scale bars: 10μm. **1F**. Proportion of *deltaA*^*pos*^ cells (cyan) within the indicated cell types (n=4 independent hemispheres, Dm). **1G**. Distribution of AAs in qNSCs and aNSC singlets according to *deltaA* expression. Red dashed line: median (n=4 independent hemispheres, Dm). Statistics: non-parametric t-test (Mann-Whitney test), p-value <0,0001.

In contrast to most epithelia, where AAs are largely homogeneous, we observed that the AA of NSCs and their aNP progeny are highly heterogeneous in shape and size (Figure 1A). This prompted us to probe for a possible significance of AA and associated parameters along lineage progression. We extracted quantitative information on various geometrical cell parameters of NSCs/NPs and asked whether they differ among cell types (NSCs vs NPs) or NSC states (qNSCs vs aNSCs). We segmented the cell contour of all progenitor cells on triple IHC of *Tg(gfap:eGFP)* fish at 3-mpf labeled with ZO1, PCNA, and *gfap*:GFP. We adapted a code previously developed for *Drosophila* epithelial tissues^39^ and quantified AA, anisotropy, perimeter length and the number of neighbors, to correlate them with molecular markers (Figures 1C and S1A). While the shapes of apical surfaces displayed a broad range of anisotropies across all cells of the germinal population, we found significant correlations of cell states and types with AA, apical perimeters, and the number of apical neighbors (Figures 1D and S1A): on average, qNSC AAs are larger (84μm^2^ ± 8,9 s.d) than those of aNSC singlets (54μm^2^ ± 17,1 s.d) and aNP AAs are even smaller (10μm^2^ ± 3,6 s.d). The small AA of aNPs fits with their delaminating behavior (Figures 1B and 1C -orange, right column-) and is the most discriminative morphological parameter compared to aNSCs (Figures S1A and B).

Two non-exclusive hypotheses could underlie the observed correlation between AA and NSC state: (i) NSC geometry could influence NSC activation, or (ii) NSCs may have distinct proliferation rates modulating their geometry (e.g., their AA). The latter hypothesis is supported below.

### *deltaA* expression correlates with cell type, state, and quantitative apical parameters

Notch signaling promotes quiescence and progenitor state maintenance in adult NSCs in zebrafish and mouse^7,32,40–42^. Thus, we also focused on this pathway as a further possible readout of NSC decisions *in vivo*. In the zebrafish adult pallium, *deltaA* expression is restricted to aNSCs and aNPs^8,32^, and aNSCs and TAPs in the adult mouse SEZ express its ortholog *Delta-like 1 (Dll1)*^20,21^.

To achieve a quantitative description, we performed quadruple IHCs on double transgenic *Tg(gfap:dTomato);Tg(deltaA:egfp)* fish to detect Tomato (*gfap*)^43^, GFP (*deltaA*)^31^, PCNA and ZO1. We considered *deltaA*^*pos*^ all cells with weak to strong IHC GFP signal, and *deltaA*^*neg*^ all cell with no visible IHC GFP signal. We found that 87% of aNSC singlets and 99% of aNPs are *deltaA*^*pos*^, while 77% of qNSCs are *deltaA*^*neg*^ (Figures 1E and 1F)^8^. Next, we found a strong anti-correlation between AA and *deltaA* expression in NSCs: *deltaA*^*pos*^ NSCs display in average a significantly smaller AA than *deltaA*^*neg*^ NSCs (p-value <0.0001), both for all NSCs and for the qNSC and aNSC states separately (Figures 1G and S1C). *deltaA*^*pos*^ qNSCs have in average smaller AAs (54μm^2^ ± 3,7 s.d) than their *deltaA*^*neg*^ counterparts (112μm^2^ ± 15,4 s.d), and *deltaA*^*pos*^ aNSCs (32μm^2^ ± 8,3 s.d) have smaller AAs than *deltaA*^*neg*^ aNSCs (91μm^2^ ± 28,2 s.d). These observations reveal global trends between *deltaA* expression, a small AA, activation status and lineage progression from NSC to NP. *deltaA* and AA may participate in or may readout lineage progression. They may be, or not, functionally interdependent factors in this process. To help better understand causal links or the respective roles of AA and *deltaA* in NSC activation, we further explored NSCs that deviate from this general trend. First, there is a broad distribution of apical parameters at the level of individual NSCs. For example, 20% of qNSCs have an AA within the 5-60μm^2^ range, corresponding to the average AA of aNSCs. Among aNSCs, 31% have an AA within the 50-150μm^2^ range, corresponding to the average AA of qNSCs (Figures 1D and S1D). Second, a measurable fraction of cells also breaks the rule linking the *deltaA*^*pos*^ and aNSC states: 23% of qNSCs are *deltaA*^*pos*^ and 13% of aNSCs are *deltaA*^*neg*^. *deltaA*^*pos*^ qNSCs have a small AA (around 50μm^2^), while *deltaA*^*neg*^ aNSCs have a large AA (above 50μm^2^) (Figure 1G and S1C). These outliers reveal that the link between *deltaA* and AA (*deltaA*^*neg*^/small AA, *deltaA*^*pos*^/large AA) is independent of NSC states.

### A novel transgenic tool and image analysis pipeline reveal NSC behavior in link with cell geometry and *deltaA* expression

To interpret the heterogeneities observed among NSCs and with NPs regarding AA and *deltaA* expression, we interrogated these parameters along NSC trajectories in real time. To retrieve fate-related cellular events (activation, division mode and delamination) together with a quantitative resolution of NSC AAs using intravital imaging, we built a new transgenic line expressing a truncated version of human ZO1^44^ fused to the mKate2 fluorescent reporter (hZO1-mKate2) under control of the *gfap* promoter^45^. Using multicolor 2-photon microscopy and double transgenic *Tg(gfap:hZO1-mKate2);Tg(deltaA:egfp)* fish crossed into the double mutant *Casper* background^46^ (Figure 2A), we performed live intravital imaging of 2.5-mpf adult fish every two to three days for at least 17 time-points (tp) (43 days). We also developed an image analysis pipeline to extract dynamic quantitative measurements from the time-lapses (Figures S2A and B). The *Tg(gfap:hZO1-mKate2)* line successfully labeled the NSC apicobasal interface with an excellent signal to noise ratio, allowing us to follow cytokinesis events -which all occur perpendicularly to the plane of the NSC layer-as well as delamination events (Figure 2B and video S1). This live analysis confirmed highly variable *deltaA:gfp* expression intensities. For simplicity, we binned the GFP signal in four intensity scores (from 0 = no expression to 3 = strong expression), validated by quantitative pixel values (Figures S2C to S2E). We also compared the values obtained for AAs in live and fixed samples and found them highly similar (R-squared 0.92, Figures S2F and S2G). As an additional calibration, to focus on divisions originating from a previously quiescent NSC, we estimated the time needed from NSC activation onset (initiation of expression of the G1 phase markers PCNA or MCM5) to cytokinesis (identified monitoring ZO1-mKate2). We imaged two 3-mpf *Tg*(*mcm5:egfp)*^29^;*Tg*(*gfap:hZO1-mKate2)* fish for over 39 days, and quantified the average activation-to-cytokinesis transition rate to be 0.2815 days^-1^ (0.2470 to 0.3235, 95% CI), i.e., in average, a transition duration of 3.5 days (Figure S3A). This fits our previous study based on *Tg*(*gfap:dTomato)*;*Tg*(*mcm5:egfp)* animals^8^. Thus, division events preceded by a minimum of 2 imaging time intervals (4 to 5 days) without division were considered to originate from a previously quiescent NSC.

**Figure 2.**
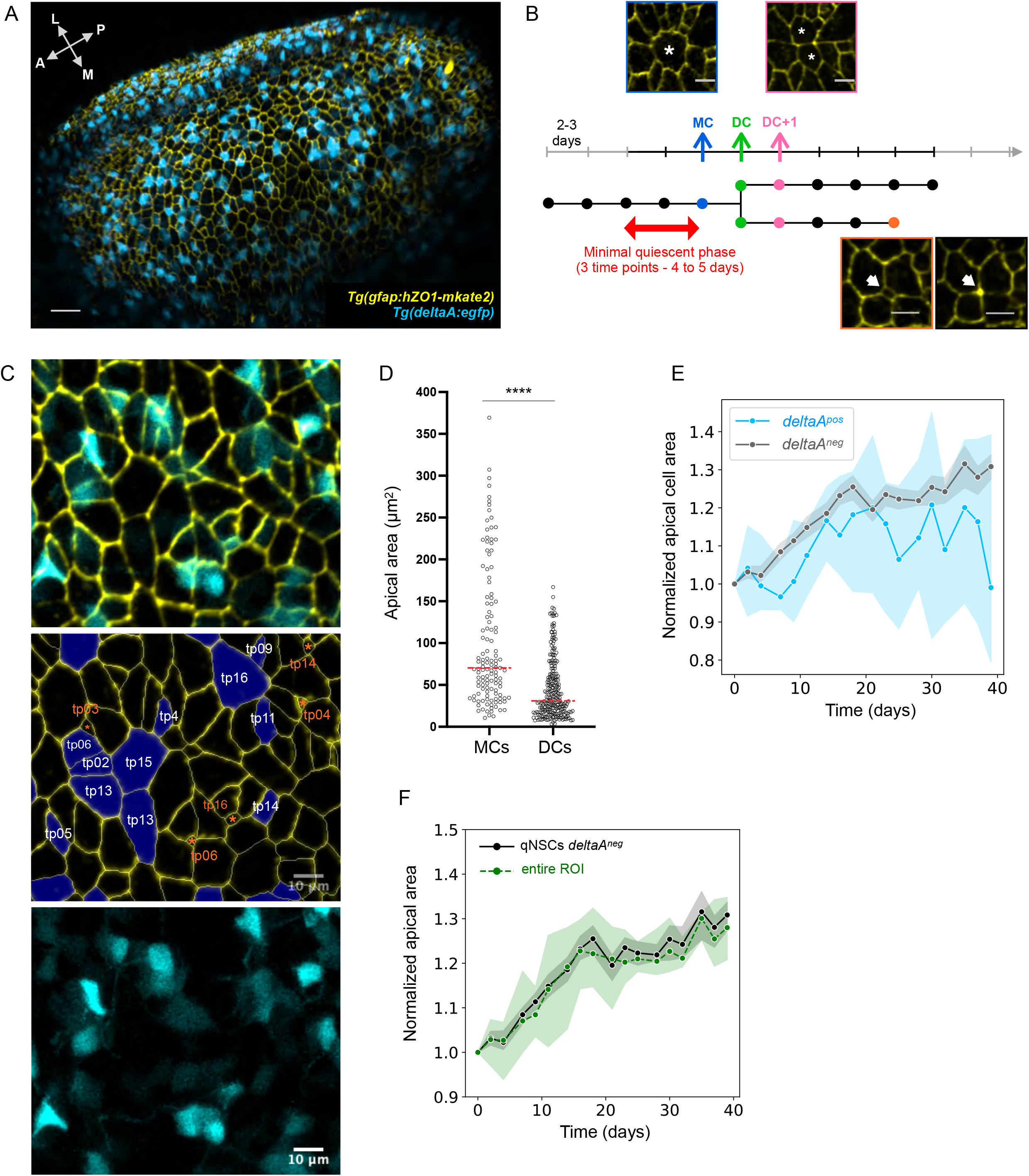
Apical area dynamics in NSCs. **2A**. Dorsal view in 3D of the pallial surface in a 2.5-mpf casper*;Tg(deltaA:eGFP);Tg(gfap:hZO1-mKate2)* fish imaged intravitally using biphoton multicolor microscopy. *deltaA*:eGFP: *deltaA* transcription, cyan; *gfap*:ZO1-mKate2: NSC apical contours, yellow. Arrows show the anteroposterior (A-P) and mediolateral (M-L) axes. Scale bar: 30μm **2B**. Schematic example of a dividing track (horizontal tree) (horizontal arrow: time; vertical bars: imaging time points (tp), consecutive tps separated by 2 or 3 days), to position a mother NSC (MC, blue arrow, example picture), Daughter NSCs (DCs, cytokinesis event, green arrow) and DC+1 at the next time point (pink arrow, example picture). DC is the first tp where two daughters can be identified. We only scored dividing tracks where MC was preceded by at least 4 to 5 days without division (black part of time arrow). The interruption of a track before the end of the recording period corresponds to a delamination (illustrated example in orange). Scale bars: 10μm. **2C**. High magnification with split and merged channels of the first tp of a time-lapse. The middle image is overlaid by a color code showing NSCs that will divide (purple, tp at which division will occur) and cells that will delaminate (orange, tp at which delamination will occur). **2D**. AA distribution of all MCs and all their DC). Red dashed line: median (n=4 independent hemispheres, Dm), 102μm^2^ for MCs and 45 μm^2^ for DCs (p-value<0.0001, Mann-Whitney test). **2E**. Normalized AA of qNSCs during non-division phases as a function of time and *deltaA* expression (*deltaA*^*neg*^ NSCs: grey line, *deltaA*^*pos*^ NSCs: blue line) (with median and 95% CI). **2F**. Normalized area of the entire region of interest over time (green line), vs normalized AA of *deltaA*^*neg*^ qNSCs during non-division phases (grey line, as in 2E) (with median and 95% CI).

Next, we fully analyzed time-lapse movies from four different *Tg(gfap:hZO1-mKate2);Tg(deltaA:egfp)* fish and could follow the fate of 828 NSCs. Most of these NSCs (634 tracks, 76.6% of all) are resting in a long quiescent phase, i.e., not dividing nor delaminating during up to 43 days of imaging. 194 cells divided at least once, enabling to explore their cell lineage. Using the criteria above, 125 of these were activation and division events from quiescence (divisions following a qNSC to aNSC transition). Finally, 97 delamination events were observed. Delaminating cells are the smallest cells tracked, with a median AA of 8μm^2^ and 98% express *deltaA*, strongly suggesting that they are aNPs (Figures S3B). They usually express *deltaA* at highest levels, confirming our fixed data analyses on the link between AA and *deltaA* expression (Figure S3C). All tracks showing a division, a delamination and/or a change in *deltaA* expression over time are shown in Figure S3D.

Hereafter, considering only divisions following a quiescence event, we will refer to the first tp after division as DC (Daughter NSC), and respectively name the preceding and following tp MC (Mother NSC) and DC+1 (one tp after the appearance of two DCs) (Figure 2B). Successive imaging tp are then referred to as DC+2, DC+3, etc. (Figure 2C). These dataset and measurements put us in a position to explore the temporal and quantitative relationship of NSC quiescence exit, division and fate with *deltaA* expression and AA dynamics.

### Cytokinesis events and slow growth during quiescence control NSC apical area dynamics

We first characterized the dynamics of NSC AAs over time to identify the events leading to AA changes. Generally, tracking individual NSCs from one tp to the next during non-division phases did not reveal significant AA modifications (Figure S3D). Unsurprisingly, the major NSC AA remodeling events are divisions, each generating two DCs of equal AA, the sum of which approximates the initial size of the MC (Figure 2D). Thus, each division leads to a decrease of AAs by half. This is expected to generate NSCs of smaller and smaller AAs through divisions, raising the question of how some NSCs reach a large AA. We therefore measured global apical expansion rates (tracking AA of non-dividing NSCs during 40 days). For non-dividing *deltaA*^*pos*^ NSCs (n=49), this rate is almost null and shows a high variability (Figure 2E). In contrast, the AAs of *deltaA*^*neg*^ NSCs (n=584) is overall growing by 30% in 40 days (Figure 2E). This affects *deltaA*^*neg*^ NSCs of small and large AA. Although growth does not appear linear (Figure 2E), it suggests a daughter could eventually regain the size of its mother in more than 100 days, a duration compatible with our previous estimations of average quiescence durations in adult pallial NSCs^14^. To further validate this observation, we reasoned that the AA growth of *deltaA*^*neg*^ NSCs should be paralleled with a global growth of the segmented and tracked area, given that *deltaA*^*neg*^ NSCs account for most NSCs overall (80%). We found that this was indeed the case (Figure 2F). Together, these results indicate that NSC AAs vary as a consequence of abrupt decreases at division, and, in the case of *deltaA*^*neg*^ NSCs, compensatory slow growth during quiescence.

### The behavior of *deltaA*^*pos*^ NSCs is biased towards proliferation and differentiation

We next assessed the overall dynamics of *deltaA* expression across time and NSC decisions. As in our static dataset (Figure 1G), we confirmed a negative correlation between *deltaA* expression and AA, and we retrieved qualitatively and quantitatively similar qNSCs and aNSCs subtypes (Figure 3A). In non-dividing tracks (n=703), *deltaA* expression appears continuous with only 2% of NSCs changing their expression profile during a time-lapse (9 NSCs switching *deltaA* OFF, 5 NSCs switching it ON). In dividing tracks (n=125), *deltaA* expression before division is also largely stable, either always ON (77 tracks) or always OFF (40 tracks), with only 9.4% of dividing NSCs changing *deltaA* level before dividing (8 tracks switch *deltaA* ON before division out of 85 dividing *deltaA*^*neg*^ MCs, and a single *deltaA*^*pos*^ track switches *deltaA* OFF) (Figure S3D). Finally, after division, when present in (a) daughter cell(s), *deltaA* expression persists in most cases for the remainder of the track (96.2 % ± 1.9 s.e.m) (Figure S3D). Thus, *deltaA* expression is not a transient state but signs a stable change of NSC signature.

**Figure 3.**
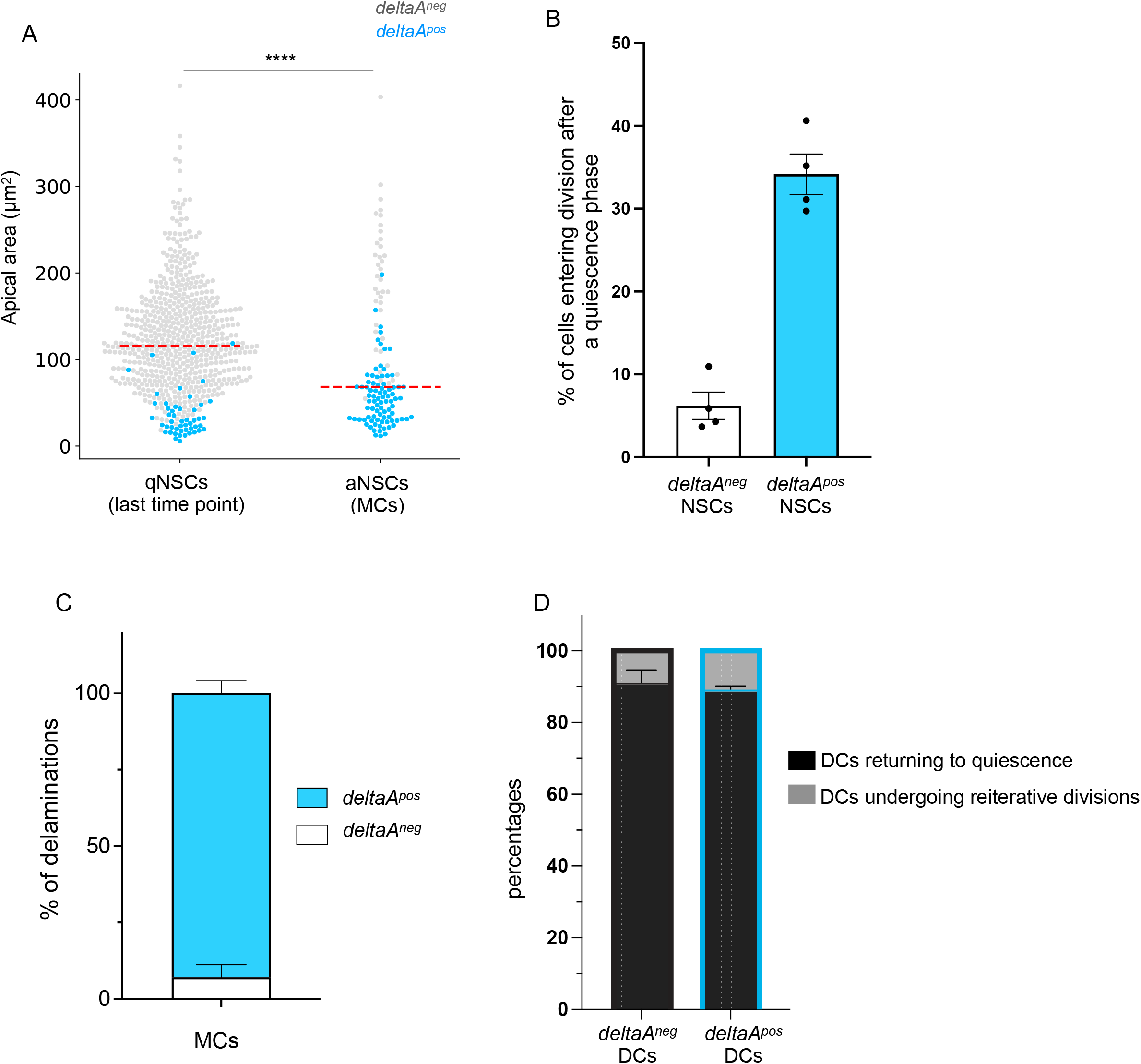
The behavior of *deltaA*^*pos*^ NSCs is biased towards proliferation and differentiation. **3A**. Distribution of AA in qNSCs (measurements on the last tp of each time-lapse) and aNSCs (measurements of MCs) according to *deltaA* expression (n=4 independent fish pooled). Red dashed line: median. Statistics: non-parametric t-test between both distributions (Mann-Whitney test, two-tailed), p-value <0.0001. **3B**. Percentages of *deltaA*^*pos*^ vs *deltaA*^*neg*^ NSC tracks where a post-quiescence division event takes place during a whole movie (n=4 fish, median with 95% CI). **3C**. Percentages of delamination events occurring in *deltaA*^*pos*^ MCs vs *deltaA*^*neg*^ MCs (n=4 fish, error bars for s.e.m). **3D**. Percentages of *deltaA*^*pos*^ vs *deltaA*^*neg*^ DCs returning to quiescence vs. undergoing reiterative divisions after a post-quiescence division (n=5 reiterative division events for *deltaA*^*neg*^ DCs, n=22 events for *deltaA*^*pos*^ DCs, error bars for s.e.m).

We then compared the proliferative and fate behavior of *deltaA*^*pos*^ and *deltaA*^*neg*^ NSCs (considering as *deltaA*^*pos*^ all NSCs where GFP is visible). First, we found that among 168 *deltaA*^*pos*^ NSCs, on average 34.2% (±2.4% s.e.m) activate from quiescence and divide during a movie, which is around 5.5 times more frequently than *deltaA*^*neg*^ NSCs (6.2% ±1.6% s.e.m, n=628) (Figure 3B). These values are confirmed using growth rates: *deltaA*^*neg*^ NSCs, in average, have a growth rate of 5.587×10^−3^ day^-1^ (doubling time = 124 days), against 2.460×10^−2^ day^-1^ (doubling time = 28 days) for *deltaA*^*pos*^ NSCs. Second, we found that the vast majority of delaminations in dividing tracks occur in the progeny of *deltaA*^*pos*^ NSCs (92.8% ± 4.1 s.d, n=45 delaminations) (Figure 3C). Third, our data also capture that the large majority of *deltaA*^*pos*^ DCs (223 over 250 DCs tracked for at least 4 days after division) re-enter quiescence post-division, a proportion similar to that of *deltaA*^*neg*^ DCs (Figure 3D).

Thus, although *deltaA* expression is neither necessary for division nor a criterion for immediate division, the fate of *deltaA*^*pos*^ NSCs is biased towards proliferation and lineage termination. This bias is visible at long term, as delamination and differentiation can occur days to weeks after *deltaA* expression onset and involve several further NSC divisions and quiescence phases. The association of *deltaA* expression to a NSC state that is engaged at long-term towards neurogenesis correlatively associates the *deltaA*^*neg*^ state with a signature of NSC stemness.

### The onset of *deltaA* expression signals the first asymmetric NSC division along the NSC lineage

The investigation of *deltaA* expression dynamics first revealed that *deltaA* expression onset tightly correlates with cytokinesis. Indeed, in all 40 division events of *deltaA*^*neg*^ NSCs, *deltaA* transcription is initiated post-division, in DCs or DCs+1 (Figure S3D). A second major observation was that dividing *deltaA*^*neg*^ NSCs systematically generate daughter cells of opposite *deltaA* expression status, one NSC daughter remaining *deltaA*^*neg*^ while the other becomes *deltaA*^*pos*^ (Figures 4A, 4C). This asymmetric outcome (ON/OFF, referred to below as “binary asymmetry”) can be already apparent for DCs and/or reinforced for DCs+1 as *deltaA* expression progressively becomes detectable in DC pairs that were initially *deltaA*^*neg*^*/deltaA*^*neg*^ at DC (Figure 4C’).

**Figure 4.**
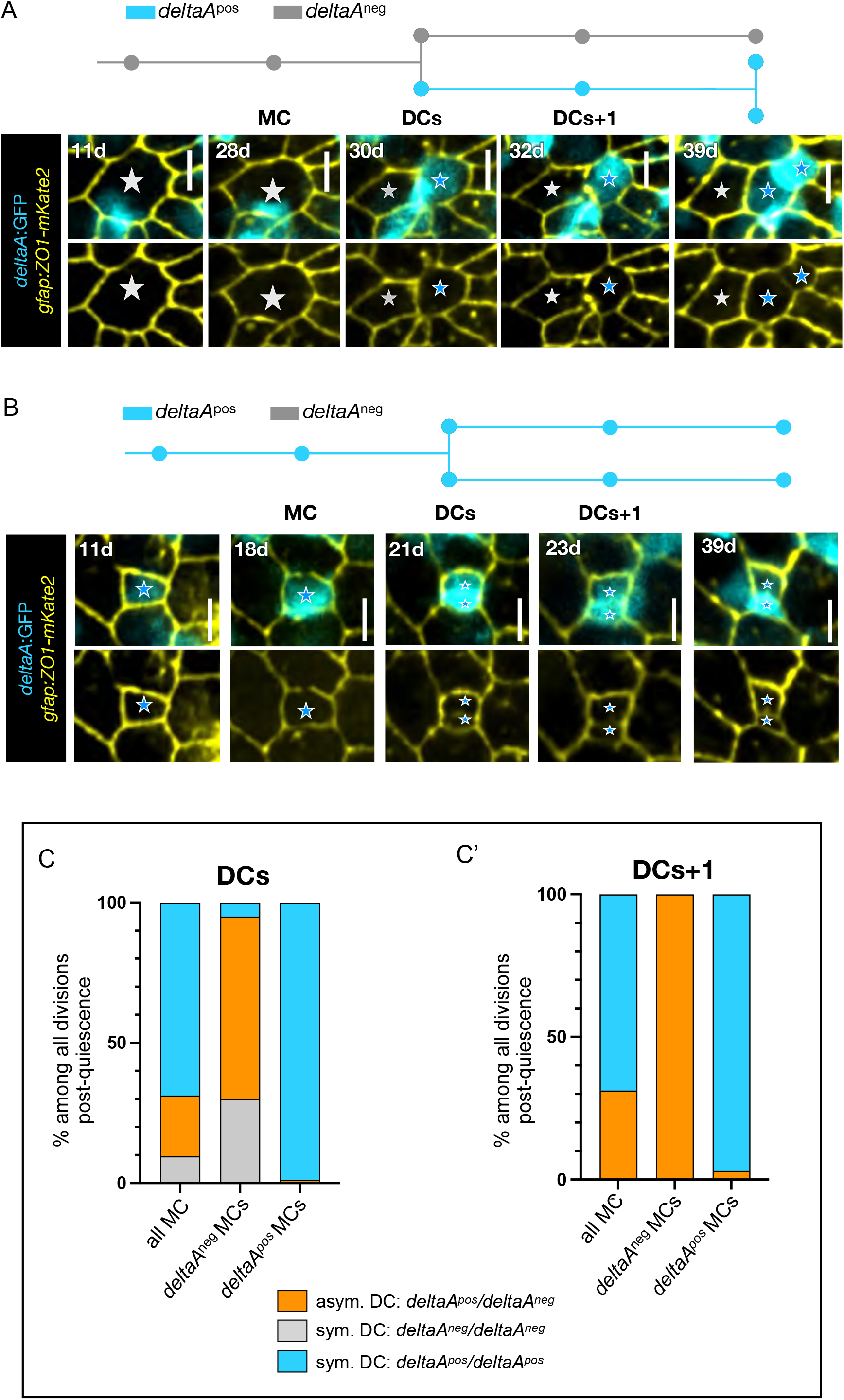
The *deltaA* expression status of NSCs predicts different division modes. **4A-4B**. Examples of asymmetric (A) and symmetric (B) divisions based on *deltaA* expression. In each case, the track (*deltaA*^*pos*^ NSCs: blue; *deltaA*^*neg*^ NSCs: grey) corresponds to the representative snapshots displayed underneath, showing *gfap*:ZO1-mKate2 (yellow) and *deltaA*:GFP (cyan) (top) or *gfap*:ZO1-mKate2 only (bottom). Stars to MC and its DCs (grey stars: *deltaA* OFF, blue stars: *deltaA* ON). **4C-C’**. Percentages of each division mode observed for DCs (C) and DCs+1 (C’) (asymmetric *deltaA*^*neg*^/*deltaA*^*pos*^: orange; symmetric *deltaA*^*neg*^/*deltaA*^*neg*^: gray; symmetric *deltaA*^*pos*^/*deltaA*^*pos*^: blue) depending on the *deltaA*^*pos*^ or *deltaA*^*neg*^ status of the MC.

In striking contrast, *deltaA*^*pos*^ NSCs systematically divided to initially generate two *deltaA*^*pos*^ daughter NSCs (Figures 4B and 4C). Ranking *deltaA* expression using intensity scores (Figures S2C to S2E) further revealed that DCs in ON/ON pairs often differ in GFP intensity (Figures S4A to S4B). A fraction of these intensity-based asymmetries transformed into ON/OFF binary asymmetries over time, in average after 11 to 12 days (Figure S4C). Finally, we asked if the anti-correlation between AA and *deltaA* expression was detectable at cytokinesis when DCs have different sizes. We found that 60% of DC pairs have AAs that differ by at least 20% and that this increases to 76% and 82% of DCs+1 and DCs+2 pairs, respectively (Figure S4D, red curve). Among such DC pairs, a higher *deltaA* expression level in the smallest DC is the predominant situation, although this is acquired over time and most DC pairs initially display equal *deltaA* levels (Figure S4D).

Together, the onset of *deltaA* expression is largely slave to a NSC division, and most divisions of NSCs activating from quiescence generate daughters of different *deltaA* expression intensities over time. The divisions of *deltaA*^*neg*^ MCs are the first asymmetric divisions of the lineage, generating a *deltaA*^*neg*^/*deltaA*^*pos*^ pair of differentially fated DCs (stemness vs neurogenesis commitment, respectively). This asymmetry is established at or immediately post-division, thus might depend on cell-cell interactions or contextual cues in DCs.

### AA and *deltaA* transcription before division are robust and independent predictors of activation propensity and binary *deltaA* asymmetry in DC pairs

AA and *deltaA* expression are strongly correlated with each other, and with NSC division frequency and division mode over time. Thus, we wondered whether these two parameters are predictors of NSC decisions, and if so, whether one parameter dominates. To first evaluate the predictive value of AA and/or *deltaA* expression pre-division on activation frequency, we conducted logistic regressions to evaluate the probability of division for qNSCs as a function of NSC AA for all tracks. In both *deltaA*^*neg*^ and *deltaA*^*pos*^ qNSCs, activation and division probability increased with AA (Figure 5A). For a given AA, this probability was higher in *deltaA*^*pos*^ NSCs (Figure 5A). Considering *deltaA* expression levels further shows that, for a given AA, high expression scores are associated with a higher division propensity than low scores (Figure S5A). However, size alone, in the absence of the *deltaA* expression parameter, is not sufficient to predict the probability of NSC division (Figures S5B).

**Figure 5.**
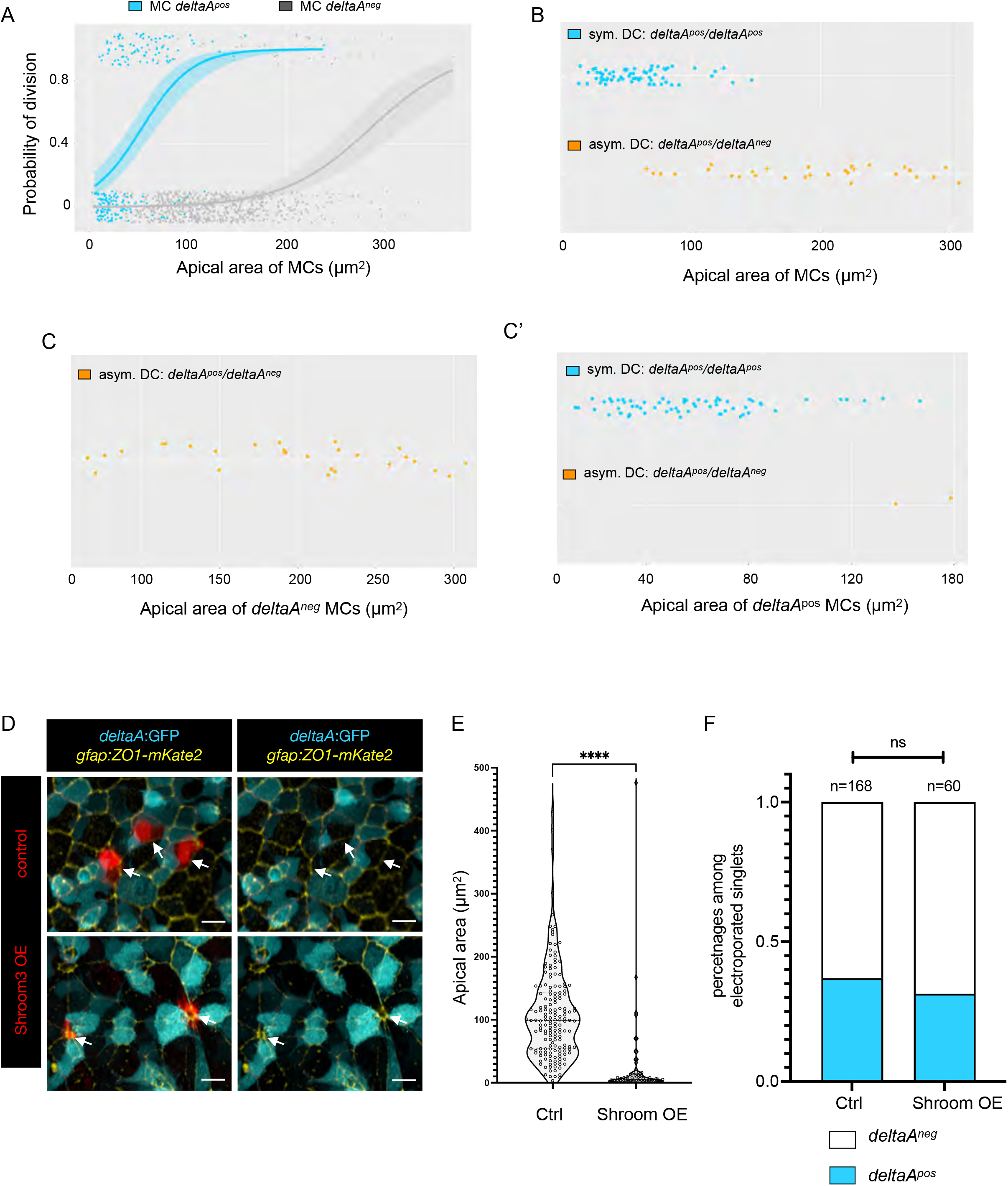
deltaA expression and AA individually predict NSC fate decisions. **5A**. Logistic regression model using AA and *deltaA* expression as covariates, showing the probability of NSC division as a function of AA for each *deltaA* expression status (*deltaA*^*neg*^ MC: gray, *deltaA*^*pos*^ MC: blue). The statistical interaction between AA and *deltaA* was found statistically significant (type II Wald χ2 tests, p-value = 4.67 ×10^−3^**). 5B-5C’**. *deltaA* expression status in DC+1 pairs (symmetric *deltaA*^*pos*^*/deltaA*^*pos*^ or asymmetric *deltaA*^*neg*^*/deltaA*^*pos*^) as a function of the apical area of MCs, (B) when all MCs are considered together (*deltaA*^*pos*^*/deltaA*^*pos*^ pair: blue, *deltaA*^*neg*^*/deltaA*^*pos*^ pair: orange) or (C) when *deltaA*^*pos*^ (blue) and (C’) when *deltaA*^*neg*^ (gray) MCs are considered separately. **5D**. High magnification of the NSC layer showing *shroom3*-electroporated NSCs (Shroom3-mCherry, bottom row; control expressing mCherry only, top row) in 3mpf *Tg(deltaA:egfp)* fish immunostained for GFP (cyan) and ZO1 (yellow). **5E-F**. Effect of Shroom3 overexpression on NSC AA (E) and the proportion of *deltaA*^*pos*^ NSCs among electroporated NSCs (F). AA: Shroom3-mCherry-overexpressing NSCs: n=70, mean AA: 14.09μm^2^; control electroporated NSCs: n=168, mean AA: 86.01μm^2^). Statistics: non-parametric t-test (Mann-Whitney test), p-value <0,0001. Proportion of *deltaA*^*pos*^ NSCs among electroporated NSCs: Shroom3-dsRed-overexpressing NSCs: n=238 NSC singlets, 31%; control NSCs: 37%). Statistics: two-sided Fisher’s exact test, p-value ns.

We next addressed the predictability power of AA and *deltaA* expression on NSC division asymmetry, focusing on the generation of *deltaA*^*neg*^/*deltaA*^*pos*^ DC+1 pairs. This binary asymmetrical outcome appears to be highly predicted by AA when all cells are considered (Figure 5B). However, size become an irrelevant parameter when *deltaA*^*neg*^ MCs are considered separately: irrespective of their AA, *deltaA*^*neg*^ mothers divide in a binary asymmetric manner (100% of cases, n=27) and *deltaA*^*pos*^ mothers generate *deltaA*^*pos*^/*deltaA*^*pos*^ DC pairs (97% of DCs+4 cases, n=66) (Figure 5C-C’).

Together, these results indicate that the non-expression of *deltaA* systematically predicts the generation of asymmetrically fated DCs upon division. It also predicts a lower division frequency overall than for NSCs expressing *deltaA*, while a large AA biases division propensity towards higher division rates both for the *deltaA*^*neg*^ and *deltaA*^*pos*^ NSC types.

### NSC AA and Notch signaling are functionally independent parameters in the control of NSC decisions along lineage progression

We next addressed whether and to which extent *deltaA* and AA may be functionally interacting parameters. To address the effect of AA on *deltaA* expression, we overexpressed the PDZ domain-containing protein Shroom3, or its dominant-negative form dnShroom3, using intracerebral injection and electroporation of Shroom3-mCherry-or dnShroom3-mCherry-encoding plasmids into pallial NSCs of *Tg(deltaA:egfp)* adults *in vivo*. These factors were reported to respectively decrease vs increase AA in embryonic epithelial cells^37^. While dnShroom3 was without effect on NSC AA in our system, Shroom3 led to extremely efficient AA shrinkage within 3 days post-electroporation, as measured using ZO1 IHC (Figures 5D and 5E). This was not associated with a significant induction of *deltaA* expression (31% in Shroom3 overexpression vs 37% in control, n=238 singlets of NSCs counted in 30 brains) (Figure 5F).

Next, we indirectly manipulated DeltaA activity by perturbing Notch signaling using the gamma-secretase inhibitor LY411575^8,47^. This treatment has the advantage of blocking signaling in both *deltaA*^*pos*^ NSCs and their neighbors. To permit a dynamic analysis with knowledge of NSC history, two of the *Casper;Tg(gfap:hZO1-mKate2);Tg(deltaA:egfp)* adult fish initially subjected to intravital imaging for 40 days, and an additional one fish imaged at this last tp, were treated with LY411575 (LY) while imaging for a further 7 days. Recording was conducted at days 1, 3, 5, 6 and 7 post-treatment onset (Figures 6A and 6B) and we reconstructed NSC trajectories (261 tracks under LY treatment) (see Figure S6A for all dividing tracks). Blocking Notch primarily activates NSCs^8,47,48^. We indeed observed a first massive wave of induced cytokinesis on the third and fifth day of treatment (Figures 6B and 6C), validating our Notch blockade procedure, here concomitant with imaging: 89 NSCs divided among 261 NSC tracked. This is 19 times more than the overall likelihood of NSCs to divide under control conditions (per cell per day, upon LY treatment: 0.0852, CI 95% is 0.0715-0.1001, vs control: 0.0044, CI 95%: 0.0038-0.0052). We found that Notch blockade was not accompanied with changes of AA during this time frame: the AA of non-dividing NSCs was stable and dividing NSCs generated DCs of half the size of their mother’s AA (Figures S6A and S6B).

**Figure 6.**
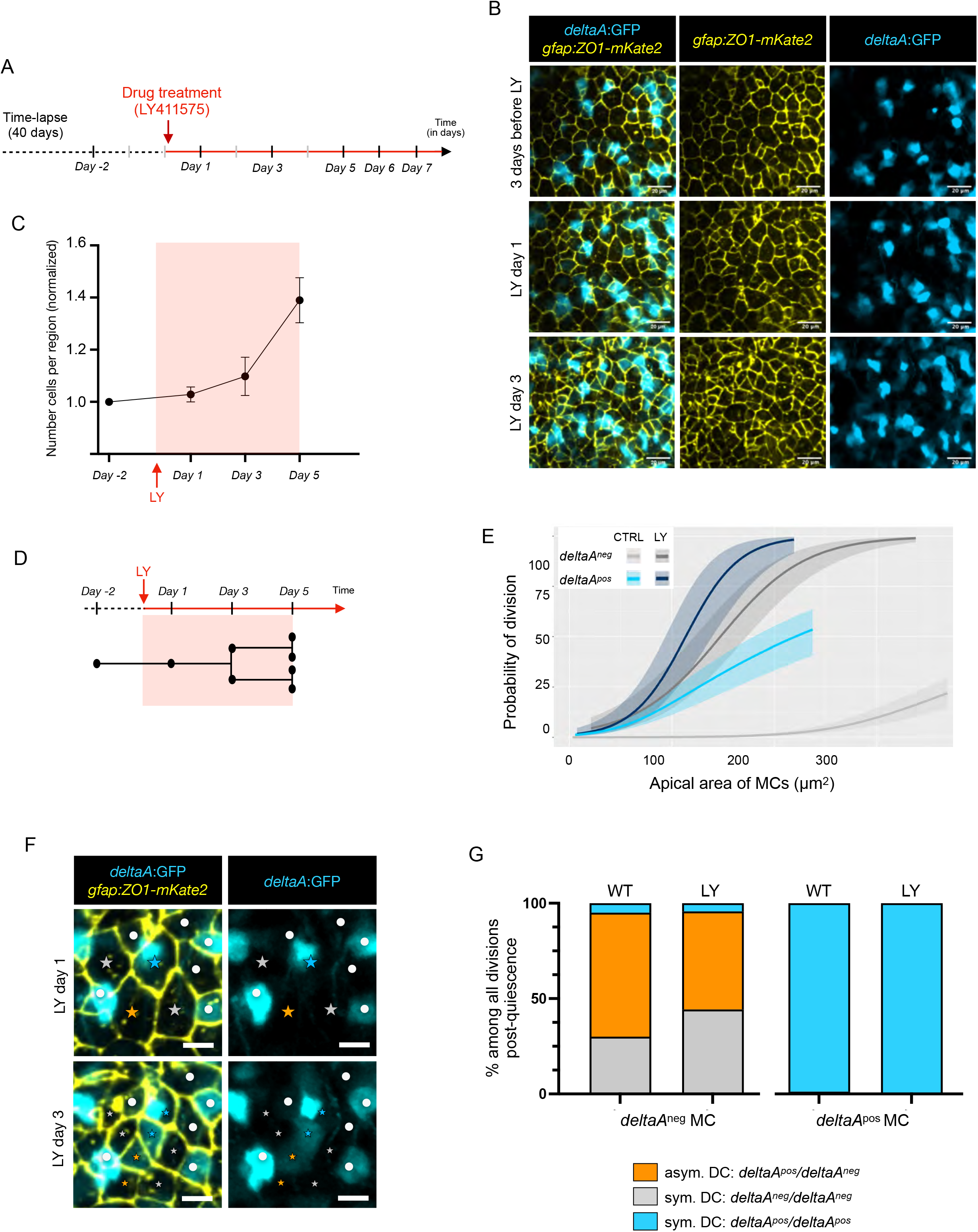
*deltaA* expression and AA in mother NSCs predict NSC decisions independently of Notch signaling. **6A**. Experimental scheme of LY treatment. LY411575 was added to the fish water for 7 days. Fish were imaged at days 1, 3, 5, 6 and 7 of treatment. **6B**. Merged and split snap-shot images of the maximum projections of a dorsal view of a live-imaged 3-mpf casper*;Tg(deltaA:eGFP);Tg(gfap:hZO1-mKate2)* fish showing progressive NSC reactivation upon LY treatment (gfap:ZO1-mKate2: yellow, *deltaA*:GFP: cyan). Scale bars: 20μm^2^. **6C**. Number of cells per segmented region (n=3 fish, mean and error bars for s.e.m), normalized to the value at day -2, as a function of time (see Figure 6A). **6D**. Time window during which NSCs could be reliably tracked upon LY treatment. **6E**. Logistic regression model using AA and *deltaA* expression as covariates: probability of NSC division as a function of AA for each *deltaA* expression status of the MC; compaired untreated (CTRL) and LY411575-treated (LY) conditions. Notch inhibition abolishes the differences in activation rate between *deltaA*^*pos*^ and *deltaA*^*neg*^ NSCs (p-value=0.16, Chi-squared test) but the AA effect persists. Predictions for control conditions: estimated by rescaling the rates over 4 days from all divisions over the duration of the movies; predictions under LY treatment: estimated using all divisions over 4 days (see Figure S6A). Individual regressions pooled for this analysis: see Figure S6C. **6F**. Merged and split time-lapse images of a live-imaged 3-mpf casper*;Tg(deltaA:eGFP);Tg(gfap:hZO1-mKate2)* fish highlighting dividing NSCs (colored stars) and non-dividing NSCs (white dots) during the first and third days of LY treatment. Colored stars and white dots: tracking of NSCs and their progeny. Scale bars: 10μm^2^. **6G**. Distribution of division modes based on the *deltaA* expression status of the DCs, for divisions having occurred after 3 days of LY treatment (n=3 fish) or in control conditions (n=4 fish).

We next used this paradigm to test whether the prediction power of AA on division propensity *in vivo* resists a context where Notch signaling is blocked. We focused on the first 5 days of treatment, as the numerous NSC division events made it difficult to faithfully connect DCs to their MC afterwards (Figure 6D). During the first 5 days of treatment, the initial expression of *deltaA:gfp* in NSCs was largely unaffected, allowing to unambiguously track both *deltaA*^*neg*^ and *deltaA*^*pos*^ NSC categories. Logistic regressions measuring NSC division frequency as a function of AA confirmed that the predictive character of AA on division propensity can be detected over a period of 4 days in control *deltaA*^*neg*^ and *deltaA*^*pos*^ NSCs (Figure 6E). Upon LY treatment, AA still appeared predictive of division propensity in both cell categories, which now displayed identical regression profiles shifted towards higher activation rates (Figures 6E and S6C). Thus, under physiological conditions, both *deltaA*^*neg*^ and *deltaA*^*pos*^ NSCs are limited in their activation frequency by ongoing Notch signaling (although at different levels or with different sensitivities). More, the predictive character of AA on division propensity operates in the absence of Notch signaling. This reveals the existence of a process biasing activation rate in a manner independent from the Notch signaling level or status of the NSC considered.

Finally, given the identified predictive power of *deltaA* expression on division mode, we addressed whether Notch signaling was involved in controlling NSC fate at division. To enrich our analysis for NSC divisions that occurred as a result of LY, we focused on DC pairs revealed after 3 days of LY treatment, i.e., with their single MC detectable at 1 day of treatment (Figure 6F). Control animals were analyzed over the same duration. We could record all three division modes, generating *deltaA*^*neg*^/*deltaA*^*neg*^, *deltaA*^*neg*^/*deltaA*^*pos*^ and *deltaA*^*pos*^/*deltaA*^*pos*^ DC pairs (Figures 6F and 6G). We specifically focused on divisions from *deltaA*^*neg*^ MCs. While their exclusive *deltaA*^*neg*^/*deltaA*^*pos*^ division fate is very rapidly acquired post-cytokinesis, around 50% of the DC pairs solidify this fate between DC and DC+1 from an initially *deltaA*^*neg*^/*deltaA*^*neg*^ fate (Figure 4C and 4D). We observed no difference in the DC fate of *deltaA*^*neg*^ MCs under Notch blockade compared to control conditions (Figure 6G) -and, as previously concluded, AA appeared irrelevant (Figure S6D). Although fate consolidation could not be studied, these results suggest that the initial steps of asymmetry generation in the NSC lineage are independent of Notch signaling. We also observed no change in the *deltaA*^*pos*^/*deltaA*^*pos*^ DC outcome of divisions from *deltaA*^*pos*^ MCs (Figure S6D).

## Discussion

Activation frequency and division modes condition NSC renewal and lineage progression, the two concomitant hallmarks of stemness. To understand how these parameters are controlled *in vivo*, we exploited long-term intravital imaging of NSC morphometric, molecular and fate readouts to identify predictors of NSC decisions in the endogenous context of the intact neurogenic niche. We identify AA and *deltaA* expression as associated although functionally unlinked parameters that correlate with NSC division propensity. We further uncover the first detectable asymmetry in the NSC lineage, where *deltaA*^*neg*^ NSCs systematically generate daughter NSCs that effectively segregate self-renewal and neurogenesis commitment. Our results provide the first NSC lineage reconstruction associated with a temporal series of predictive cellular and molecular hallmarks in situ. These hallmarks underscore the relevance of intrinsic cues associated with the *deltaA*^*neg*^ status in the accomplishment of stemness, a conclusion reinforced by our demonstration that these cues are independent of Notch signaling.

### A model of NSC lineage progression based on apical surface area and *deltaA* expression

The transgenic line *Tg(gfap:ZO1-mKate2)* reveals the dynamics of NSC apical membranes, now allowing to investigate multiple individual cell or collective features such as apical cell geometries or neighborhoods, over different numbers of cells, in link with cell decisions (such as cytokinesis or delamination) and molecular markers (here *deltaA*). Tracking for at least 40 days the behavior of >800 NSCs in their endogenous niche inside the adult zebrafish pallium^8,29^, we could reconstruct a presumptive lineage spanning months of an NSC’s life. Focusing on cell-intrinsic features, we show that AA and *deltaA* expression, together, sign NSC position along lineage progression (Figure 7): (i) the divisions of *deltaA*^*neg*^ NSCs generate *deltaA*^*pos*^ NSCs, placing *deltaA*^*neg*^ NSCs hierarchically upstream, (ii) NSC AAs grow slowly over time together with an increasing probability to divide, (iii) conversely, high *deltaA* levels and a small AA sign temporal proximity to NSC pool exit and neuronal differentiation. Importantly, in between these extremes, we identify a key lineage transition linked to both parameters and impacting fate: the division of *deltaA*^*neg*^ NSCs. This event systematically generates differently fated daughter NSCs, of which the *deltaA*^*neg*^ daughter has the potential to regrow its AA and behave identically to its mother, while the *deltaA*^*pos*^ daughter does not regrow, and engages into more frequent subsequent divisions and ultimately differentiation. Altogether, these results, supported by quantitative measures of division rates and AA growth, allow inferring relative temporal and hierarchical NSC states from in vivo images and interpreting static datasets staining NSCs in their niche.

**Figure 7.**
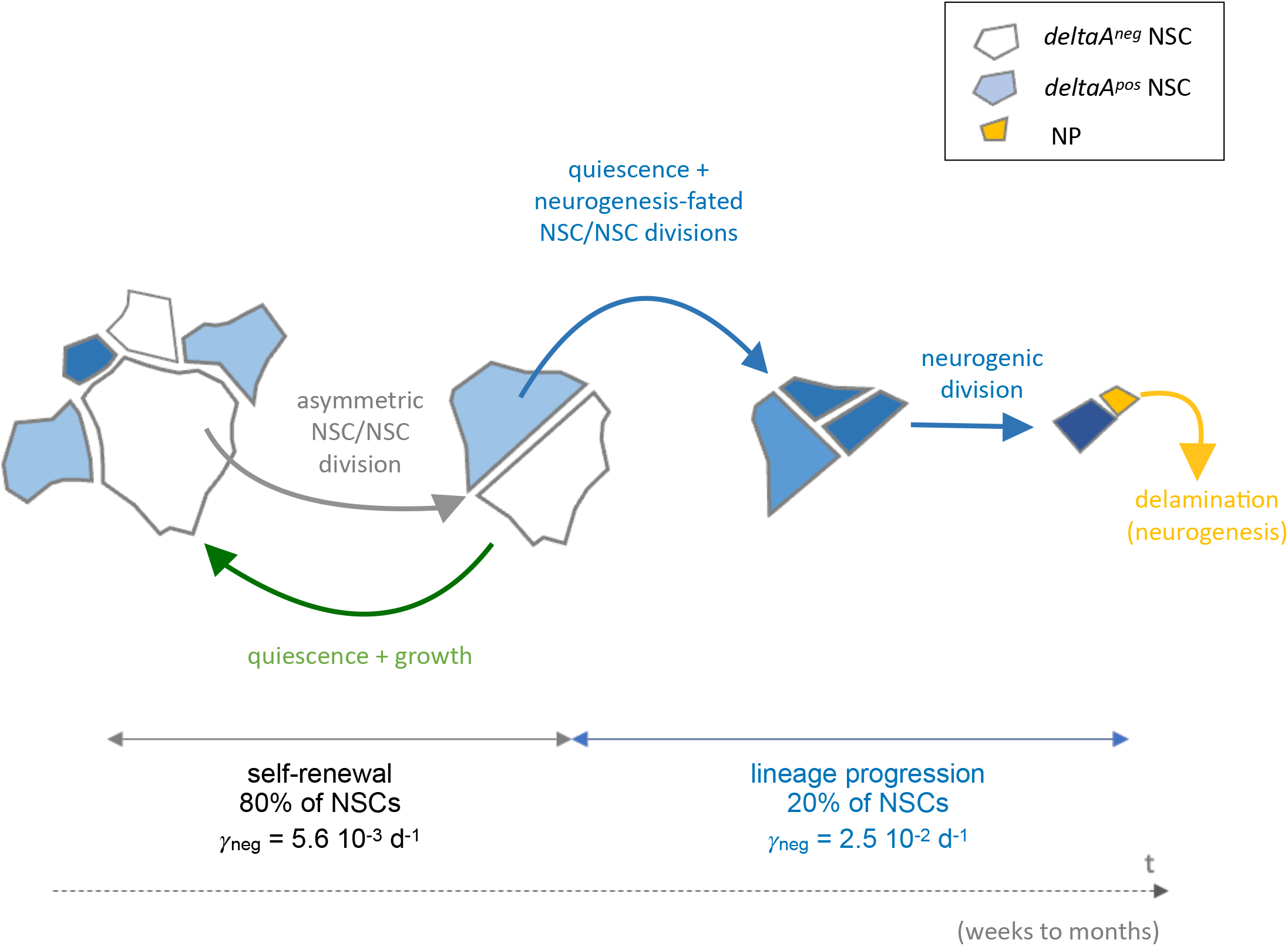
Dynamic hierarchy of self-renewal and lineage progression based on AA and *delta* expression in adult pallial NSCs in situ. Schematic apical representations of NSCs and NPs, with relative AAs and *deltaA* expression (color coded), in an interpretative drawing resulting from the assembly of overlapping tracks, covering a time frame of weeks to months. The division of *deltaA*^*neg*^ NSCs signs the transition from self-renewal to neurogenesis commitment. γ_neg_ and γ_pos_ are average activation rates for *deltaA*^*neg*^ and *deltaA*^*pos*^ NSCs respectively.

### AA reads out activation propensity in adult NSCs

Our results point to AA as a cellular readout correlated with NSC division propensity. In embryonic radial glia, AA varies dynamically with nuclear position and cell cycle phase^49^. The adult pallium is very different however, as there is no interkinetic nuclear migration, and NSC AA during quiescence is either stable (in *deltaA*^*neg*^ NSCs) or growing at a steady rate over weeks (in *deltaA*^*pos*^ NSCs) (Figure 2E). We also did not notice AA increase pre-division (Figure S3D). Our work does not attempt to solve the mechanisms leading to AA increase, although the observed growth rate suggests an active process rather than a passive phenomenon e.g., due to stretching to accommodate size changes in neighbors.

In the embryonic retina, experimentally enlarging AA via dnShroom3 favors the proliferating over differentiating neural progenitor fate, in a process mediated by enhanced Notch signaling^37^. Overexpression of dnShroom in adult NSCs *in vivo* did not enlarge AA, while, as observed in embryos, Shroom overexpression triggered massive AA reduction and delamination, precluding to directly probe AA impact on division frequency. Whichever mechanisms link AA and division propensity, our current results argue that this mechanism is Notch-independent, since we did not observe an induction of *deltaA* expression in Shroom3-overexpressing NSCs (Figures 5D to 5F), and the link between AA and division propensity persists even upon LY treatment (Figure 6E).

AA may be *per se* informative or read out another correlated geometry or sub-cellular organization feature (Figure S1). NSCs also possess a large basolateral component, which branches over several hundreds of microns in the parenchyma and could receive or encode division-related signals^50,51^. AA however appears as a hub that at least reads out pertinent information relative to NSC division propensity, within a given *deltaA* expression status. In *deltaA*^*neg*^ NSCs, AA growth between divisions implies that AA positively correlates with time in quiescence, which might be one of the measured parameters. There is, however, no threshold below or above which *deltaA*^*neg*^ NSCs will systematically divide or remain quiescent, and further mechanistic work is needed to interpret this correlation. In the case of *deltaA*^*pos*^ cells, 100% of cells with an AA <10um^2^ delaminate during a 43 day-movie (Figure S3F). A hierarchical relationship between AA decrease and fate acquisition remains to be studied, but Shroom3 overexpression indicate that delamination can also be induced by AA decrease in the absence of a fate change (as read by *deltaA*).

Finally, our dynamic intravital imaging results shed light on the mechanisms that account for the correlation between AA and *deltaA* expression in the adult pallial NSC population. At any time, static images show that *deltaA*^*neg*^ NSCs are generally large and *deltaA*^*pos*^ NSCs generally small (Figure 1). This is highly reminiscent of the checkerboard pattern modeled in embryonic neuroepithelia and interpreted to result from dictating Notch signaling directionality by the surface of contact between signaling and receiving cells^52^. In the adult NSC population, where cell divisions and AA changes occur, our results support a distinct interpretation where AA differences between *deltaA*^*neg*^ and *deltaA*^*pos*^ NSCs emerge from two parameters: a regulation of *deltaA* transcription onset by lineage progression, coupled with the lower division frequency of *deltaA*^*neg*^ NSCs and their AA regrowth during quiescence.

### The restriction of NSC potential is progressive, and stemness is signed by the *deltaA*^*neg*^ status

Clonal tracing and intravital imaging revealed that adult NSCs can divide according to three possible division modes (NSC/NSC, NSC/NP or NP/NP) in mouse and zebrafish^8–15^. This classification is based on the generation of the NP terminal fate, and it is a major question to understand whether all NSCs are equal along lineage progression until this fate decision^12,14,17^. Our results highlight asymmetric and overall generally increasing *deltaA*:GFP levels at each division of a *deltaA*^*pos*^ mother (Figure S3C). Although these observations await confirmation by monitoring DeltaA protein -we currently lack an antibody detecting DeltaA in adult zebrafish NSCs-, they suggest that sister *deltaA*^*pos*^ cells are generally not equivalent, and that fate acquisition is a progressive process along the division sequence of each *deltaA*^*pos*^ NSC. This is highly reminiscent of the progressive transition from commitment to differentiation described in mouse skin stem cells^53^. There was a trend but no clear-cut AA-related rule associated with the assignment of *deltaA*:GFP differences between daughters (Figure S4D). In addition, the massive induction of proliferation by LY treatment precluded analyzing *deltaA*:GFP levels between *deltaA*^*pos*^ daughters in the absence of Notch, and the mechanisms involved in progressive fate restriction within the *deltaA*^*pos*^ lineage remain open.

Importantly, our work also reveals that *deltaA*^*neg*^ NSCs systematically generate daughters of opposite *deltaA* status. While the *deltaA*^*pos*^ NSC engages towards a neurogenic fate at long term, several arguments suggest that the *deltaA*^*neg*^ daughter behaves identically to its mother: it will never turn on *deltaA* expression (Figure S3D), it has the potential to regrow to the initial mother size (Figure 2E), and all *deltaA*^*neg*^ NSCs, whatever their size, follow this asymmetric division mode (Figures 4C and 4C’). These observations show that the *deltaA*^*neg*^ daughter is engaged in self-renewal and identify this NSC/NSC division as the first asymmetric division of the NSC lineage, generating two NSC daughters of different potential that segregate stemness maintenance from neurogenesis. The systematic outcome of *deltaA*^*neg*^ NSC divisions likely implies a cell-autonomous process and raises the question of its control. The *deltaA* status is tracked using the *Tg(deltaA:gfp)* transgene^54^, implying transcriptional regulation of *deltaA* expression post-division. Asymmetric segregation of the DeltaA protein itself could be directly monitored in embryonic neural progenitors *in vivo*^55,56^, and in adult NSCs overexpressing Dll1-GFP *in vitro*^20^. This is unlikely to drive the first NSC/NSC asymmetry identified here for the pallial lineage, since we do not detect *deltaA*:GFP expression prior to division. Ascertaining this point will nevertheless require the direct detection of DeltaA. Finally, we found that the *deltaA* expression asymmetry is initially insensitive to Notch blockade (Figure 6G). This argues against a mechanism such as intra-lineage regulation, involving Notch-mediated sister-sister interactions that can occur downstream of the asymmetric segregation of Notch pathway regulators other than Delta^57–59^.

### Integration of NSC heterogeneity modalities

Heterogeneities in NSC potential are postulated based on single-cell transcriptomics, BrdU or genetic fate tracing, and intravital imaging^5^. The temporal interpretation of the former approaches is inferred from statistical analyses at successive time points, as individual NSCs are not tracked over time. Complementarily, intravital imaging allows direct longitudinal tracking but generally does not read gene expression to infer molecular progression^8,13,15,60^. It now remains crucial, but a challenge, to integrate these different modalities into a comprehensive understanding of NSC behavior at the individual cell and population levels. In particular, it is unresolved whether self-renewal originates from stochastic fate decisions within the main lineage^27^, identifies a sub-lineage within the NSC population^13,60^, or is an upstream state in a NSC hierarchy^14,17^. Supporting the latter hypothesis, recent mathematical models in the adult mouse hippocampus^17^ and zebrafish pallium^14^ predicted a hierarchical organization of NSCs into sub-populations of different dynamics and fate, where a reservoir/dormant NSC population feeds into a more active operational/resting population responsible for neuronal production. In the zebrafish, quantitative predictions of population size, activation rates and division modes could further be inferred from the clonal data^14^. Specifically: the reservoir was postulated to account for 61% of all NSCs, to display an activation rate γ_r_ = 0.007 days (doubling time 97 days), and to be engaged in asymmetric NSC/NSC divisions generating one reservoir and one operational NSC. The operational pool was predicted to account for the remaining 39% of NSCs and, with γ_o_ = 0.023 days (doubling time 30 days), to stochastically choose between the NSC/NSC, NSC/NP and NP/NP division modes with a bias towards neurogenesis. Strinkingly, the lineage progression model that we propose here qualitatively and quantitatively fits these predictions when *deltaA* negativity vs expression is used to sign the reservoir vs operational pools, respectively: *deltaA*^*neg*^ NSCs make 80% of the total, and divide in an asymmetric *deltaA*^*neg*^/*deltaA*^*pos*^ NSC/NSC fashion with an average activation rate γ_neg_ = 0.0056 days (doubling time 124 days), while *deltaA*^*pos*^ NSCs (20% of NSCs) display NSC/NSC, NSC/NP or NP/NP division modes (Figure S3D) with an overall average activation rate γ_pos_ = 0.0246 days (doubling time 28 days) and a final neurogenic output. These comparable cell behaviors and figures, of similar orders of magnitude, solidify both sets of data towards a hierarchical organization of NSC dynamics. Together, these results stress the invaluable contribution of our approach to directly overlay, for the first time, three modalities of NSC heterogeneities: mathematical predictions, gene expression changes, and lineage progression. Such inclusive approaches set the stage for a comprehensive multi-modal understanding of NSC population dynamics *in vivo*.

## Acknowledgements

We thank the ZEN team for constant input, Lucas Sancerre for his help with Matlab and Alexis Villars for help with linear regression analyses. Funding: Work in the L. B-C. laboratory was funded by the ANR (Labex Revive), La Ligue Nationale Contre le Cancer, CNRS, Institut Pasteur, DIM ELICIT (with E.B.) and the European Research Council (ERC) (AdG 322936). L.M. was recipient of a PhD student fellowship from Labex Revive and the Fondation pour la Recherche Médicale (FRM). M.L. was recipient of a 4^th^ year PhD fellowship from the FRM. Multiphoton equipment in E.B.’s laboratory was partly supported by the ANR (ANR-11-EQPX-0029 Morphoscope2, ANR-10-INSB-04 France BioImaging). Work in Y.B.’s lab was supported by CNRS and the Institut Curie.

## Authors contribution

L.M. generated and analyzed the static dataset. L.M. generated the *Tg(gfap:ZO1-mKate2)* line with help from S.O and conducted intravital imaging, with advice from N.D, P.M. and E.B. provided help with imaging; E.M and S.B. provided expert fish care; B.G. and Y.B. provided training and support with segmentation and dynamic image analysis algorithms; V.B. and C.B. performed some analyses and plotting with Python; F.C. performed statistical analyses and regressions with R; M.L. conducted the Shroom3 overexpression experiments; S.H., MS.P. and JY. T. wrote codes for image analysis with Fiji and Python; N.D. and L.B-C. designed the study; L.M., N.D. and L.B-C. wrote the manuscript, with input from all authors.

## STAR * METHODS

### KEY RESOURCES TABLE

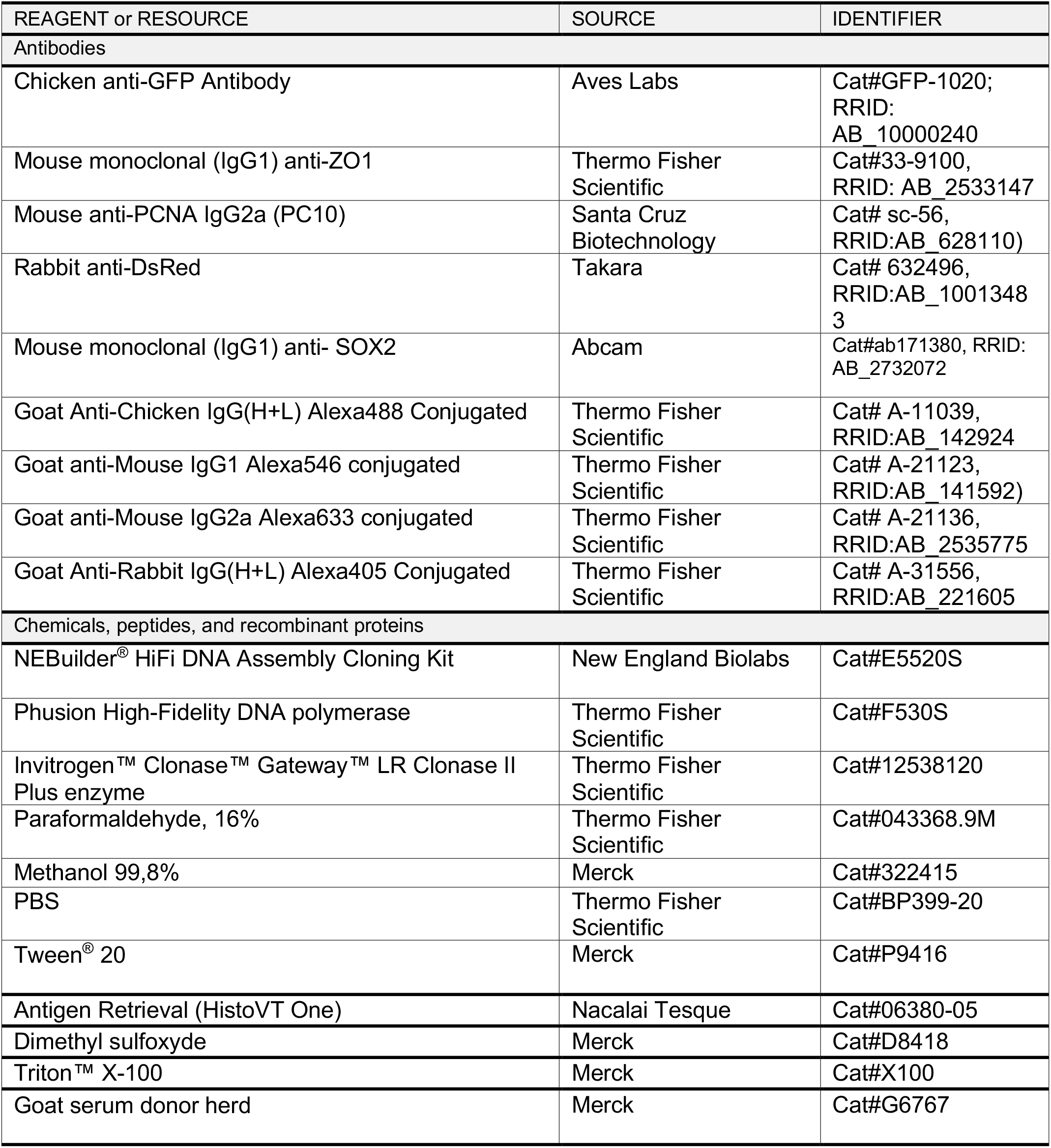
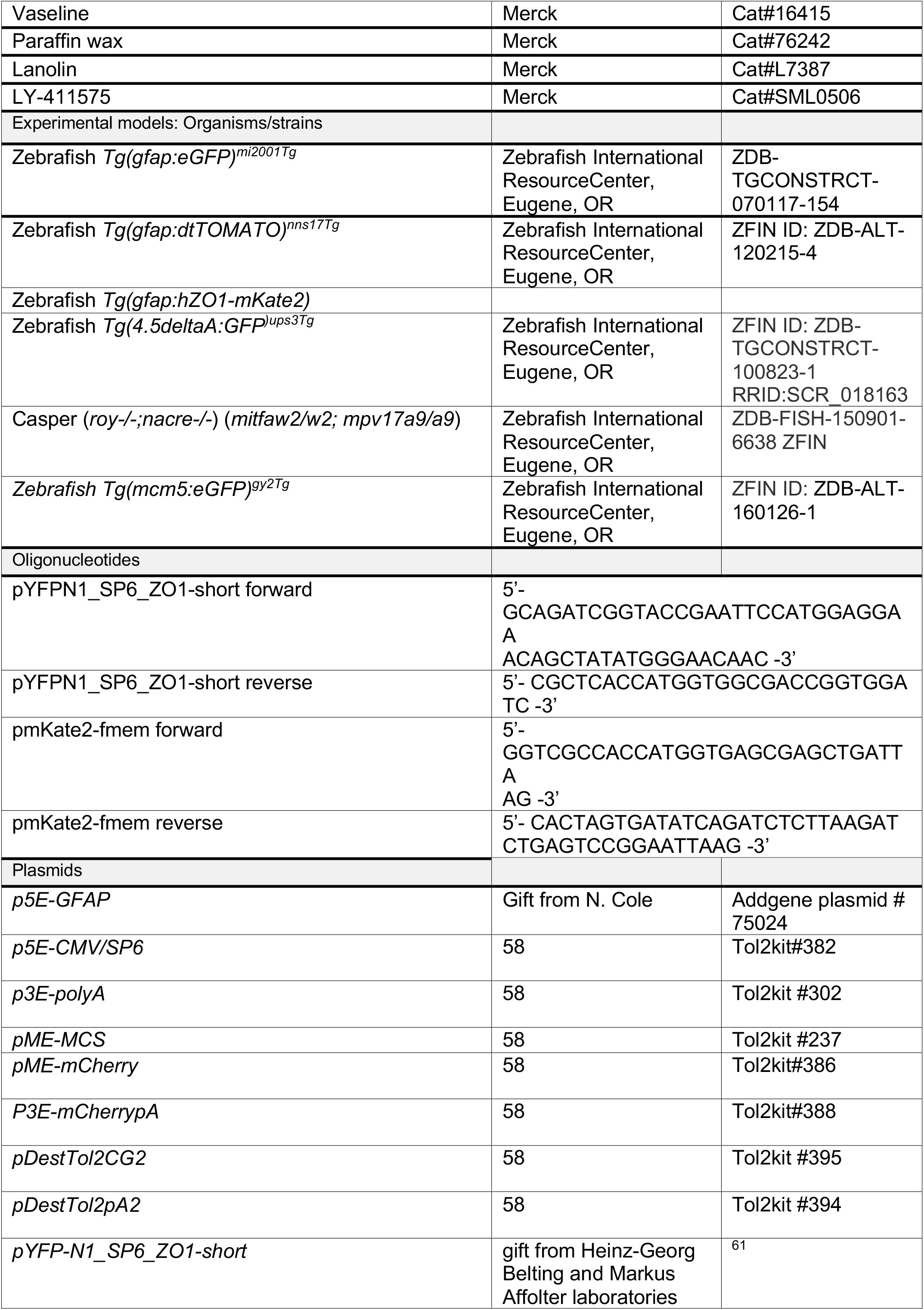
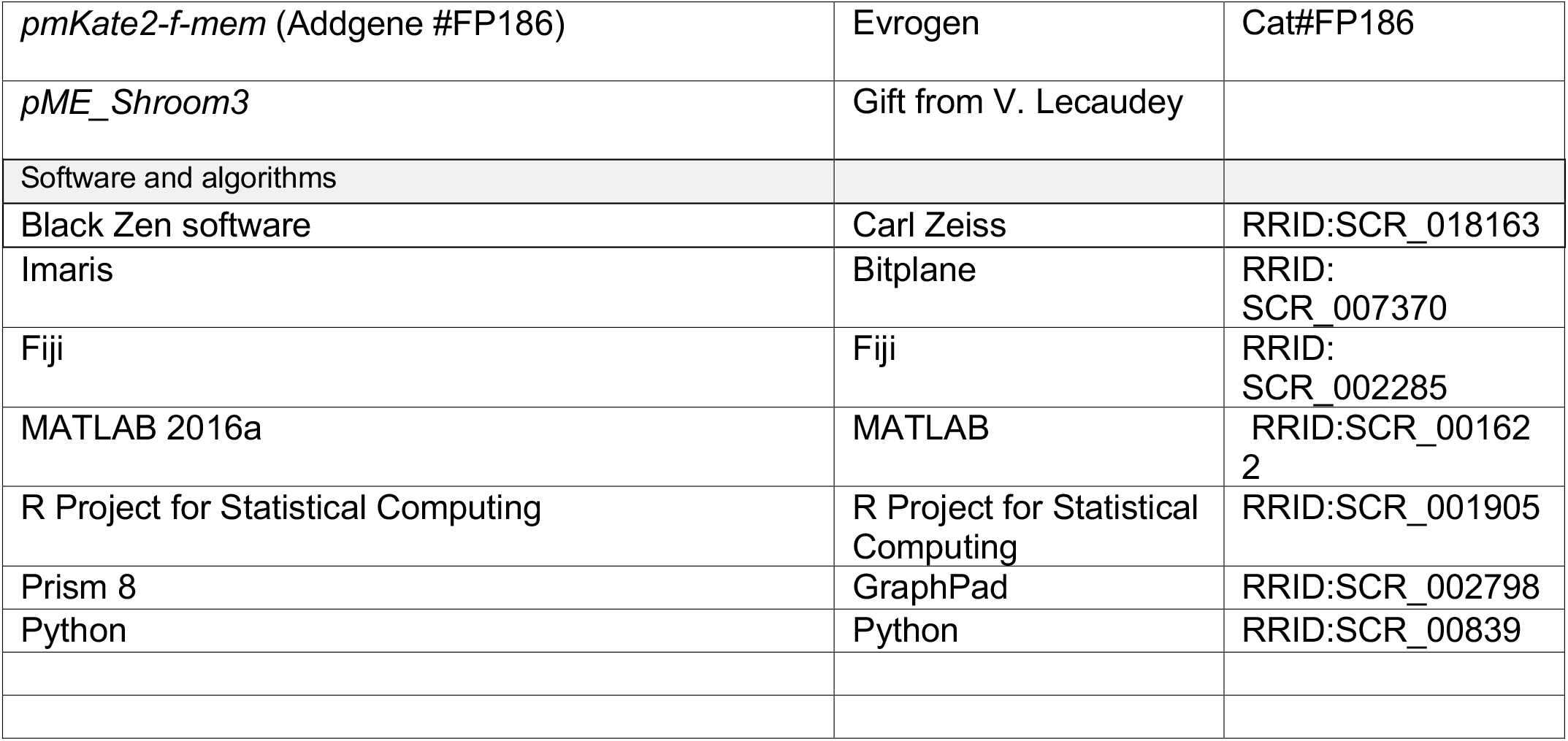

### LEAD CONTACT

Further information and requests for resources and reagents should be directed to and will be fulfilled by the Lead Contact, Nicolas Dray.

### DATA AND CODE AVAILABILITY

Codes, tabular data and main raw data are available <u>upon</u> request.

### EXPERIMENTAL MODEL AND SUBJECT DETAILS

#### Fish husbandry and lines

All animal experiments were carried out in accordance to the official regulatory standards of the department of Paris (agreement numbers C75-15-22 and A91-4772 to L.B.-C., N.D. and L.M.) and conformed to French and European ethical and animal welfare directives (project authorization from the Ministère de l’Enseignement Supérieur, de la Recherche et de l’Innovation to L.B.-C.).

All procedures relating to zebrafish (*Danio rerio*) care and treatment are conformed to the directive 2010/63/EU of the European Parliament and the council of the European Union. Zebrafish were kept in 3.5-liter tanks at 28.5°C and pH 7.4 water and in salinity-controlled conditions. They were maintained on a 14 hours light / 10 hours dark cycle and fed three times a day with rotifers until fourteen days post-fertilization (14 dpf) and with standard commercial dry food (Gemma Micro from Skretting*) afterward. All transgenic lines – Tg*(gfap:eGFP)* ^45^; Tg*(gfap:dtTOMATO*) ^43^; Tg(*mcm5:eGFP*)^gy2^ (Dray et al., 2015), Tg*(4*.*5deltaA:GFP)* ^31^ and Tg*(gfap:hZO1-mKate2)* (this paper) were maintained in the *Casper* double mutant background (*roy-/-*; *nacre -/-*) ^46^. Heterozygosity of each transgene was respected upon crosses except for the Tg*(gfap:hZO1-mKate2*) line that is visible in an adult only with two copies or more (we have multiple insertion lines). Ages of the fish are explicitly stated in the respective experiments. All fish were euthanized in ice-cold water (temperature below 4°C) for ten minutes.

### METHOD DETAILS

#### Generation of the transgenic line *Tg(gfap:hZO1-mKate2)*

The *hZO1short-linker-mKate2* sequence was assembled using the NEBuilder^®^ HiFi DNA Assembly Cloning Kit (NEB), optimized for a 3-fragments reaction following the manufacturer’s instructions. *hZO1short* (encoding human ZO1 protein lacking its actin-binding domain) followed by a linker and *mKate2* ^62^ fragments were cloned by PCR from plasmid *pYFP-N1_SP6_ZO1-short* (gift from Heinz-Georg Belting’s and Markus Affolter’s laboratories) and *pmKate2-f-mem* (Evrogen), using the Phusion High-Fidelity DNA polymerase (ThermoFisher Scientific). To generate the transgenic line, the multisite Gateway technology (Thermo Fisher Scientific) was employed, taking advantage of Tol2kit plasmids ^63^. *hZO1-mKate2* was first recloned into the *pME-MCS* plasmid (Tol2kit #237) by EcoRI and BcuI digestions, and then recombined with *p5E-GFAP*, and *p3E-polyA* (Tol2kit # 302) in *pDestTol2CG2* backbone (Tol2kit #395, containing the transgenesis marker *cmlc2:egfp-polyA*). We performed microinjection of the constructs into 1-cell *Casper* embryos together with 40 ng/μl of *transposase* capped mRNA. F0 adults crossed with *Casper* fish were screened for transmission of fluorescence (cardiac GFP and mKate2) in F1 embryos in order to generate a stable line. We chose to work with a multiple insertion line based on the quality and intensity of the signal obtained on adult fish with a 2-photon microscope.

#### Time-lapses

Anesthesia was initiated by soaking the fish for 2 to 5 minutes in water containing 0.02% MS222 (Merck). They were then transferred into a water solution of 0.005% (v/v) MS222 and 0.005% (v/v) isoflurane to maintain the anesthesia during the whole duration of the imaging session ^29^. Overall, fish were anesthetized for about 30 minutes per session and the recovery time (in freshwater without any drugs) after a session was less than 5 minutes.

#### Immunohistochemistry (IHC)

Brains were dissected in 1X solution of phosphate buffered saline at a temperature of 4°C and directly transferred to a 4% paraformaldehyde solution in PBS for fixation. They were fixed for 2 to 4 hours at room temperature under permanent agitation. After four washing steps in PBS, brains were dehydrated through 5 to 10 minutes series of 25%, 50% and 75% methanol diluted in 0.1% tween-20 (Merck) PBS solution and kept in 100% methanol (Merck) at −20°C. Rehydration was performed using the same solutions, and then brains were processed for whole-mount immunohistochemistry (IHC). After rehydration, the telencephali were dissected out and subjected to an antigen retrieval step using Histo-VT One (Nacalai Tesque) for 1 hour at 65°C. Brains were rinsed three times for at least ten minutes in a 0.1% DMSO and 0.1% Triton X-100 (Merck) PBS 1X solution (PBT) and then blocked with 4% normal goat serum in PBT (blocking buffer) 4 hours at RT on an agitator. The blocking buffer was later replaced by the primary antibody’s solution (diluted in blocking buffer), and the brains were kept overnight at 4°C on a rocking platform. The next day, brains were rinsed five to ten times over 24 hours at room temperature with PBT and incubated in a solution of secondary antibodies diluted in PBT overnight, in the dark, and at 4°C on a rocking platform. After three rinses in PBT over 4 hours, brains were transferred into PBS. Dissected telencephali were mounted in PBS on slides using a 0.7 mm-thick holders. The slides were sealed using Valap, which is a mixture of Vaseline (Merck), paraffin (Merck), and lanolin (Merck).

Primary antibodies were used at a final concentration of 1:500 for GFAP and PCNA, 1:250 for DsRed and 1:200 for Sox2 and ZO1. Secondary antibodies were all used at a final concentration of 1:1000.

#### Image acquisition

Images of whole-mount immunostained telencephali were acquired on a confocal microscope (LSM700 Zeiss) using a 40X oil objective. We acquired images with a z-step of 0,65 μm. We averaged each line four times with an image resolution of 1024×1024 pixels with a bit-depth of 12-bits. The power of the lasers was kept constants for all the acquisitions and the GAIN (the voltage of the photomultipliers) was adjusted for each experiment. We recorded mosaics with a 15% overlap to image an entire hemisphere per fish.

Intravital imaging was performed on a dual-beam 2-photon microscope (TriM Scope II, LaVision BioTec) with a 25x, 1.05 NA water immersion objective (Olympus). The mKate2 fluorophore (Tg(*gfap:hZO1-mKate2*)) was sequentially excited at 1120 nm with an optical parametric oscillator (80 MHz, ∼100-200fs pulses after the objective, Insight DS+ from Spectra-Physics) and the GFP fluorophore (Tg(*deltaA:eGFP*)) was excited at 950 nm with a titanium:sapphire oscillator (80 MHz, ∼100-150 fs, Mai Tai HP from Spectra-Physics). The mean powers after the objectives were about 30-40 mW at 1120 nm and 9 mW at 950 nm. Emitted photons were splitted with a dichroic mirror (Di02-R561-25×36, Semrock) and detected with two GaAsP detectors (H7422-40, Hamamatsu). Each line was averaged two times and acquired sequentially for each laser to avoid crosstalk at the emission (pixel dwell time of 2.42 μs). Images were acquired with a field of view of 520×520 μm and spanning a depth range of 150-200 μm (voxel size of 0.29 by 0.29 by 2 μm). The initial plane of imaging was located approximately 150 μm underneath the skin surface.

#### Pharmacological treatment

Inhibition of Notch signaling was performed using the LY411575 gamma-secretase inhibitor (Merck). The stock solution of LY411575 at 100 μM in DMSO was prepared and stored at −20ºC.

For treatment, LY411575 was applied in the fish swimming water at a final concentration of 10 μM ^747^. The solution was renewed every 24h. Control fish were treated with the same final concentration (0.1%) of DMSO carrier.

#### Overexpression of Shroom3

To generate the constructs used in overexpression experiments, the multisite Gateway technology was employed, taking advantage of Tol2kit plasmids ^63^. To build the Shroom3-mCherry plasmid, *p5E-CMV/SP6* (Tol2kit#382), *pME-Shroom3* and *p3E-mCherrypA* (Tol2kit#388) plasmids were recombined in pDestTol2pA2 (Tol2kit#394). For the negative control, *p5E-CMV/SP6* (Tol2kit#382), *pME-mCherry* (Tol2kit#386) and *p3E-pA* (Tol2kit#302) plasmids were recombined in pDestTol2pA2 (Tol2kit#394).

#### Ventricular Microinjections and Electroporations

Micro-injections into the adult pallial ventricle were performed on anesthetized fish as described^64^. For plasmid electroporations, plasmid DNA was diluted to a final concentration of 500 ng/μL in 1 x PBS and injected into the ventricle. Fish were then administered four electric pulses (50 V, 50 ms width, 1,000 ms space). Fish were sacrificed three, five, and fourteen days after the injections.

#### Processing and analysis of IHC images

All of our analyses were performed on images with 3 channels: one channel with the AA staining (ZO1), one channel with nuclear staining (PCNA), and one channel with cytoplasmic staining (either *deltaA* or gfap). Using Fiji, we performed for all channels a 3D median filter, a max Z projection, and a substract background (rolling ball radius of 50 pixels). To automatically segment AA staining with a low rate of error we performed a flat field correction on this channel only to efficiently set a homogeneous intensity (our channel of interest was divided by the duplicated version of the same channel modified with a sizeable gaussian blur >30). In the same purpose and if needed, we also performed an enhanced local contrast filter (CLAHE, blocksize = 20). Cell contours were determined, and individual cells were identified using a seed-based region growing algorithm, followed by several rounds of manual corrections. Nuclear and cytoplasmic stainings were detected in single cells using the 20 to 30% highest pixel values (Pixel values were ordered from least to most significant and the range of the 20% highest value was selected using a given percentile as a threshold) followed by manual corrections. All these analyses are done with a custom Matlab script Y. Belaïche).

#### Processing and analysis of time-lapses images

We mounted the movies composed of z-stacks with two channels (ZO1 and *deltaA*) taken on different days using a custom Fiji macro. We performed translational and rotational spatial registration manually using the correct drift function on Imaris software. Then we used the CSBDeep toolbox to denoise the apical membrane channel (ZO1). CSBDeep is a content-aware image restoration (CARE) that requires to be trained with a set of high resolution and low-resolution images. CARE is an open-source Python algorithm ^65^ and it has been adapted and trained with images from the notum of the Drosophila embryo. The restored channel, together with the second channel (*deltaA*) are MAX projected in 2D. To improve the tracking we first applied slight deformations over the image frames using the Image J’s plugin bUnwarpJ^66^. The information of all image channels was used to compute the transformation matrix between two consecutive frames. We chose the middle frame as the reference and aligned all the other frames from the referenced frame. The transformation matrices were saved in text files and served as registering other movies and performing the revert transform. The segmentation and tracking of the cell contours and *deltaA* from the registered image frames were done in Tissue Analyzer^67^. We then aligned the segmented apical surfaces back to the pre-registration state in order to compute their original apical areas. The revert transform of the image frame was deduced from its transformation file. We provide the scripts that wrap up all the steps. Dividing and non-dividing tracks are processed separately, and further analyses are performed on R and PRISM (tabular dataset are also provided).

### QUANTIFICATION AND STATISTICAL ANALYSIS

The same experimenter carried out all the segmentation and corrections for a given experiment. Balanced ratios of females and males were included in the different experimental groups as much as possible.

For the IHC and intravital dataset, means and medians are per hemisphere per animal. The size of the analyzed regions differs from fish to fish. Thus we collected different numbers of tracks per fish (data available on Zenodo). Statistical analyses were carried out using PRISM, R and Python. The normality of the residuals of the responses was assessed using normality probability plots and the homogeneity of the variance was inspected on a predicted versus residual plot. Non-parametric tests were used when the responses show a deviation of the residuals from a normal distribution and/or heterogeneous variances. In that case, overall effects were assessed with a Kruskal-Wallis test and all pairwise comparisons with the Mann-Whitney-Wilcoxon test comparing if two independent samples were selected from populations having the same distribution. All statistical analyses performed were two-tailed and their significance level was set at 5% (α=0.05).

#### Logistic regressions

##### Data sources and modelling process

Each NSC track was labelled individually according to whether the NSC underwent division during the study period. For NSCs that divided, their AA and *deltaA* expression prior to the division timepoint were recorded, while for NSCs that did not divide, the average AA and most prevalent *deltaA* expression (off or on) over the study period were used. For NSCs that divided, the *deltaA* expression of the two daughter cells was also tracked, with three possible values: *deltaA*^*neg*^/*deltaA*^*pos*^ (for an asymmetric division), *deltaA*^*pos*^/*deltaA*^*pos*^, or *deltaA*^*neg*^/*deltaA*^*neg*^.

Estimation of the proliferative capacity and fate of the NSCs was carried out using logistic regression models, with AA and *deltaA* expression as covariates. To account for heterogeneity between fish and between treatments (WT *vs* LY), the models were adjusted by adding fixed effects to the intercept and testing for interactions with other covariates. The identifiability of these fixed effects was ensured by the inclusion of two fish in both treatments. Inclusion and exclusion of parameters in the models were performed according to type II Wald χ2 tests.

##### Model for the proliferative capacities of NSCs

1106 individual tracks (844 from WT, 262 from LY) were used in this model, accounting for 214 divisions (125 from WT, 89 from LY). Both *deltaA* expression and AA, as well as an interaction term between these two covariates, were statistically significant in predicting NSC proliferative capacity. The model also had to be adjusted with an interaction term between treatments and *deltaA* expression.

As tracks were recorded over different time periods for the two treatments (around 35 days for WT, 4 days for LY), the probabilities of division estimated by the models correspond to a NSC tracked for the corresponding lengths of time. In order to directly compare the two estimates, we rescaled the estimate for WT to the shorter 4-day time period by assuming memorylessness of the NSC division process in WT fish: in this case, noting *p*_*k*_ the probability of a NSC dividing over a time period of *k* days, we can relate the probability of not dividing for *k* days, (1 − *p*_*k*_), with that of not dividing for 1 day, (1 – *p*_*1*_),

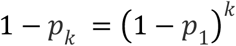

which yields a simple transformation to scale the probabilities to the same time frame,

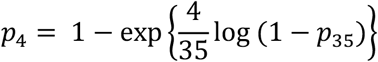

##### Model for the proliferative fate of NSCs (MCs)

As NSCs expressing *deltaA* almost exclusively yielded *deltaA*^*pos*^/*deltaA*^*pos*^ divisions (83 *deltaA*^*pos*^/*deltaA*^*pos*^ vs 1 *deltaA*^*neg*^/*deltaA*^*pos*^ in WT, and all *deltaA*^*pos*^/*deltaA*^*pos*^ in LY), only NSCs not expressing *deltaA* were using in the fit of the model. This accounted for 110 NSC divisions (41 from WT, 69 from LY). As there are 3 modalities for the *deltaA* expression of the daughter cells (*deltaA*^*neg*^/*deltaA*^*pos*^, *deltaA*^*pos*^/*deltaA*^*pos*^, *deltaA*^*neg*^/*deltaA*^*neg*^), a multinomial log-linear regression model, rather than a logistic, was fitted. Type II Wald tests revealed no parameter other than a fixed effect on the intercept accounting for the heterogeneity between fish was statistically significant in predicting the proliferate fate of the NSCs.

