## supplementary legends and figures for "Apical size and *deltaA* expression predict adult neural stem cell decisions along lineage progression"

#### Supplementary Figure Legends

**Supplementary figure S1 (related to Figure 1). Apical area and *deltaA* expression by cell type and state.**

**S1A.** Boxplots showing the distribution of cell anisotropies, perimeters, AAs and numbers of neighbors as a function of cell types and states (qNSCs, aNSCs and aNPs). n=4 independent hemispheres, Dm region, same samples as in Figure 1G.

**S1B-1C.** Average cell AA for each brain (same as in Figures 1D and 1G) as a function of cell types and states (B) and in relation with *deltaA* expression (C).

**S1D.** Histograms showing the distribution of AA of qNSCs, aNSCs and aNPs localised in the dorsal medial (Dm) region of whole-mount pallia of 3-mpf Tg(*gfap:GFP*) fish immunostained with GFP, PCNA, ZO1 and SOX2 (n=4 fish, bin size: 10 $\mu m^2$). A two-tailed non-parametric ANOVA test was performed to assess the similarity of the 3 distributions (Kruskal-Wallis test, p-value = 0,9241).

**Supplementary figure S2 (related to Figure 2). Imaging and analysis pipeline to quantify the dynamics of NSC apical geometry *in situ*.**

**S2A.** Pipeline for movie analyses. See Materials and Methods section for a precise description of each step. The images shown are examples of pre- and post-processing for each image treatment.

**S2B.** 2D projection methods. Four different projection methods were tested on the time-lapses to project in 2D the ZO1-mKate2 signal: Maximum projection and local Z projector (both available on Fiji), preMosa (available source code on the preMosa GitHub) and CARE (python code available on the CSBdeep GitHub). Zoom on a very low-resolution area in the image to compare the four projection methods. We selected CARE as the best method to resolve the ZO1 staining and improve the segmentation afterwards.

**S2C.** Closer look at *deltaA*:GFP expression intensities by color-coding eGFP intensity (FIRE lookup tables). Manual correction of the *deltaA* signal is necessary to ensure a correct assignment of *deltaA* expression to individual NSCs, because the apical surface and the corresponding underlying cell cytoplasm, which is GFP-positive, are not always in perfect register. Thus, many NSCs negative for *deltaA* are wrongly classified as positive if neighboring a balloon-shaped *deltaA*$^{pos}$ cell. Green arrows show examples where the GFP signal from one cell invades a neighbouring AA in 2D.

**S2D.** Manual scoring of the *deltaA*:GFP signal to add quantitative information to the segmentation of each cell, based on the fire LUTs and using visual and temporal criteria (see Materials & Methods). Examples of NSCs with no *deltaA* (0), weak *deltaA* (1), medium *deltaA* (2), and strong *deltaA* (3) levels. For analyses using quantitative *deltaA* values (Figures S3C, S4B and S5A), *deltaA* expression was calculated using the manual segmentation. The value of *deltaA*:GFP expression for NSCs classified as "no *deltaA*" is set at 0 and the values for other NSCs are calculated as the sum of the pixel intensity normalized by the apical area.

**S2E.** Comparison of the exact quantitation of *deltaA*:GFP expression (normalized by the area of each cell, see 2D) with manually assessed intensity scores for all cells in one time-lapse.

**S2F.** Alignment of live and fixed images. We are able to trace back previously live-imaged NSC apical surfaces on 3-mpf double transgenic *Tg(gfap:hZO1-mKate2);Tg(mcm5:gfp)* fish after fixation and IHC for SOX2, ZO1 and PCNA. Left: live-imaged tissue; right: fixed and immunostained tissue. The images on the right panels show the segmentation of the same group of cells.

**S2G.** Comparison of AAs in live and fixed/immunostained samples. Scatter plot showing AA before fixation as a function of AA after fixation for the same cell. Linear regression line with 95% CI (slope =

0,89 and R squared = 0,92). The average AA ratio between fixed AA and live AA is calculated as follows: AA ratio fixed/live = (Fixed AA - Live-imaged AA) / (Live-imaged AA) x 100. In average, fixed AA are 5% larger than live-imaged AA. The statistical difference between the two segmented regions was assessed by a two-tailed non-parametric t-test (Mann-Whitney) (n.s. p-value = 0,6667) (n = 95 cells on 2 fish).

**Supplementary figure S3 (related to Figure 2). Dynamic analysis of NSC fates in dividing tracks from intravital imaging data.**

**S3A.** Estimation of the aNSC transition rate from activation to cytokinesis, measured from intravital imaging data of *Tg(mcm5:egfp);Tg(gfap:ZO1-mKate2)* fish. The fraction of remaining aNSCs (*mcm5^pos^*) before division obtained from 137 dividing tracks (circles and bars are the average and STD from 2 fish over about 40 days) was fitted to a decaying exponential curve (red). The best fit for the transition rate is $\gamma$ = 0.28 $day$-1 with (95% CI is 0.2470 to 0.3235).

**S3B.** Apical area of cells at the imaging time point preceding a delamination event (boxplots with all datapoints, n=97). Blue dots: *deltaA*$^{pos}$ cells (representing 98% of the events). It is to note that aNPs are indirectly identified in the *Tg(gfap:ZO1-mKate2);Tg(deltaA:egfp)* background; they are *deltaA*$^{pos}$ and delimited by the ZO1-positive interface of neighbouring NSCs.

**S3C.** Representation showing the AA (in $\mu m^2$) as a function of ranked *deltaA* intensity scores. The size of the dots is proportional to AA and the colours highlight *deltaA* expression (normalized by the area of each cell). We found an inverse correlation between *deltaA* expression and AA similar to our IHC data, validating our manual segmentation.

**S3D.** Representation of all dividing tracks (n= 194, from 828 NSCs tracked in 3 fish), classified according to *deltaA* expression of MCs, and showing *deltaA* expression (*deltaA*$^{pos}$ NSCs: blue, *deltaA*$^{neg}$ NSCs: black) and AA (y axis, in $\mu m^2$) as a function of time (x axis, in days) (each dot is an imaging time point). Red arrowheads mark the different DCs for each track (division events post-quiescence phases, note that there can be several such events along the same track when DCs enter quiescence before dividing again). Green arrowheads mark the DCs from reiterative divisions (successive division events that occur without a quiescence phase in between).

**Supplementary figure S4 (related to Figure 4). Dynamics of *deltaA*:GFP intensity in daughter cells from *deltaA*$^{pos}$ NSC mothers**

**S4A.** Representative snapshot of a *deltaA*$^{pos}$ MC generating *deltaA*$^{pos}$/*deltaA*$^{pos}$ DCs of asymmetric *deltaA*:GFP intensities.

**S4B.** Proportion of *deltaA*$^{pos}$/*deltaA*$^{pos}$ DC pairs displaying an "equal" *deltaA* intensity (blue) (DCs from an individual MC have the same score for GFP intensity) vs a "unequal" intensity (orange) (DCs have different scores), following division of a *deltaA*$^{pos}$ MC.

**S4C.** *deltaA* expression at long term in DC pairs from *deltaA*$^{pos}$ MCs, showing that some *deltaA*$^{pos}$/*deltaA*$^{pos}$ DC pairs resolve into *deltaA*$^{pos}$/*deltaA*$^{neg}$ pairs (orange) over time (n=5 cell pairs among 85 pairs tracked over >12 days) (note that we progressively track fewer DC pairs due to delaminations or re-iteratives divisions).

**S4D.** *deltaA*:GFP intensity over time within DC pairs originating from a *deltaA*$^{pos}$ MC. Red curve: percentage of DC pairs where DCs differ in AA by at least 20%, for DC, DCs+1 and DCs+2. Over time, almost all DC pairs become asymmetric in size. Bar plots: distribution of the relative expression intensities of *deltaA*:GFP (color coded) within DC pairs of AA differing by at least 20%.

**Supplementary figure S5 (related to Figure 5). *deltaA* expression levels dominate the AA parameter in predicting the probability of NSC activation.**

**S5A.** Apical area and level of *deltaA*:eGFP expression together strongly predict activation of NSCs as seen using logistic regression models using both as covariate. Regressions using the four levels of *deltaA*:eGFP expression (Figure S2D) showing the trend to more activation when *deltaA*:eGFP expression increases.

**S5B.** Logisitic regression model showing the probability of NSC division as a function of AA, showing no predictability (p-value=0.454). Comparison between this model and the one using both AA and *deltaA* (Figure 5A) shows strong and significant improvement for the latter model (p-value $<2x10^{-16}$, likelihood test).

**Supplementary figure S6 (related to Figure 6). Dynamic analysis of NSC fates in dividing tracks from intravital imaging data upon LY411575 treatment.**

**S6A.** Representation of all dividing tracks (n= 89, from 261 NSCs tracked in 3 fish) upon LY treatment, classified per fish (named outi, moji and soli) and showing *deltaA* expression ($deltaA^{pos}$ NSCs: blue, $deltaA^{neg}$ NSCs: black) and AA (y axis, in $\mu m^2$) as a function of time (x axis, in days) (each dot is an imaging time point, 0 is the first day of LY treatment). Red arrowheads mark the DCs of each track (division events post-quiescence phases).

**S6B.** Normalized AA of NSCs that divide during LY411575 treatment showing that their apical area is not affected upon Notch blockade (black curve : mean of their AA normalized over the AA of MCs, considering the sum of both DCs for the tp post-division; 96% CI are indicated) (n=69 dividing NSCs over 3 fish).

**S6C.** Logistic regression model using AA and *deltaA* expression as covariates, showing the probability of NSC division as a function of AA for each *deltaA* expression status of the MC and comparing untreated (CTRL) and LY411575-treated (LY) conditions, with individual fish (names at the top of each column) shown separately (see Figure 6E for the global analysis). Regressions for control conditions (bottom row) are done by calculating the average rates over 4 days from all divisions over 40 days. Regressions under LY treatment conditions (top row) are done using all divisions over 4 days.

**S6D.** Comparison of DC fates, depending on the *deltaA* expression status of the MC ($deltaA^{neg}$ vs $deltaA^{pos}$), as a function of AA, in LY vs control treatment conditions. Each dot represents a DC pair. There is no visible effect of Notch inhibition on division modes, as read by the expression of *deltaA*.

#### Supplementary video

**Video S1 (related to Figures 2, S2, 3, S3, 4, S4, 5A-C and S5). Example of an intravital imaging dataset over 39 days.** 2D projection of a 3D-stack of the pallial surface in a 2.5-mpf casper*;Tg(deltaA:eGFP);Tg(gfap:hZO1-mKate2)* fish imaged intravitally using 2-photon microscopy. *deltaA*:eGFP: *deltaA* transcription, cyan; *gfap*:ZO1-mKate2: NSC apical contours, yellow. Some division events are highlighted with white arrowheads ($deltaA^{neg}$ NSCs) and cyan arrowheads ($deltaA^{pos}$ NSCs).

### Figure S1

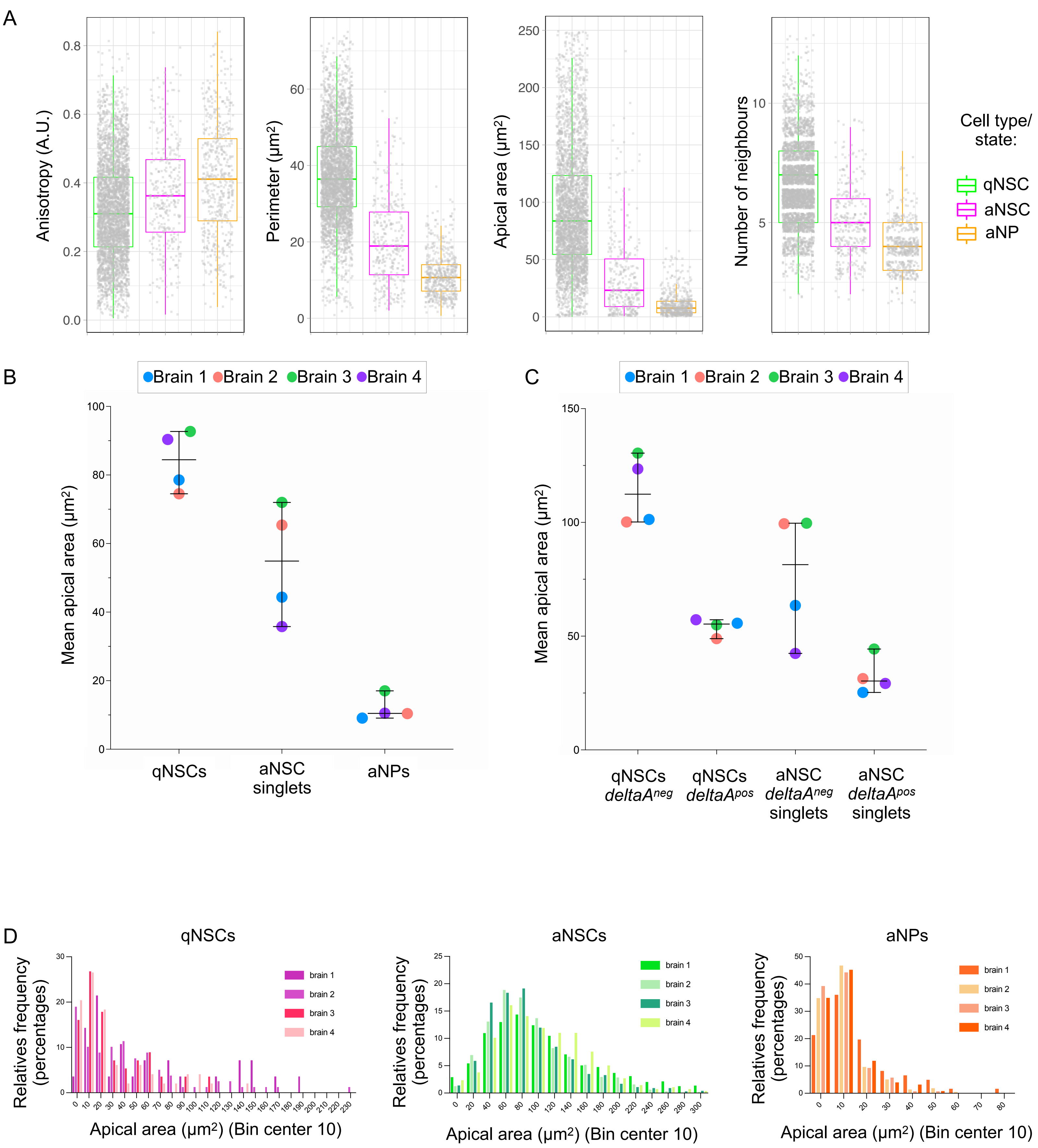

A
Anisotropy (A.U.)
Perimeter (µm²)
Apical area (µm²)
Number of neighbours
Cell type/ state:
qNSC
aNSC
aNP
B
Brain 1 Brain 2 Brain 3 Brain 4
Mean apical area (µm²)
qNSCs
aNSC singlets
aNPs
C
Brain 1 Brain 2 Brain 3 Brain 4
Mean apical area (µm²)
qNSCs deltaAneg
qNSCs deltaApos
aNSC deltaAneg singlets
aNSC deltaApos singlets
D
qNSCs
aNSCs
aNPs
Relatives frequency (percentages)
brain 1
brain 2
brain 3
brain 4
Apical area (µm²) (Bin center 10)

### Figure S2

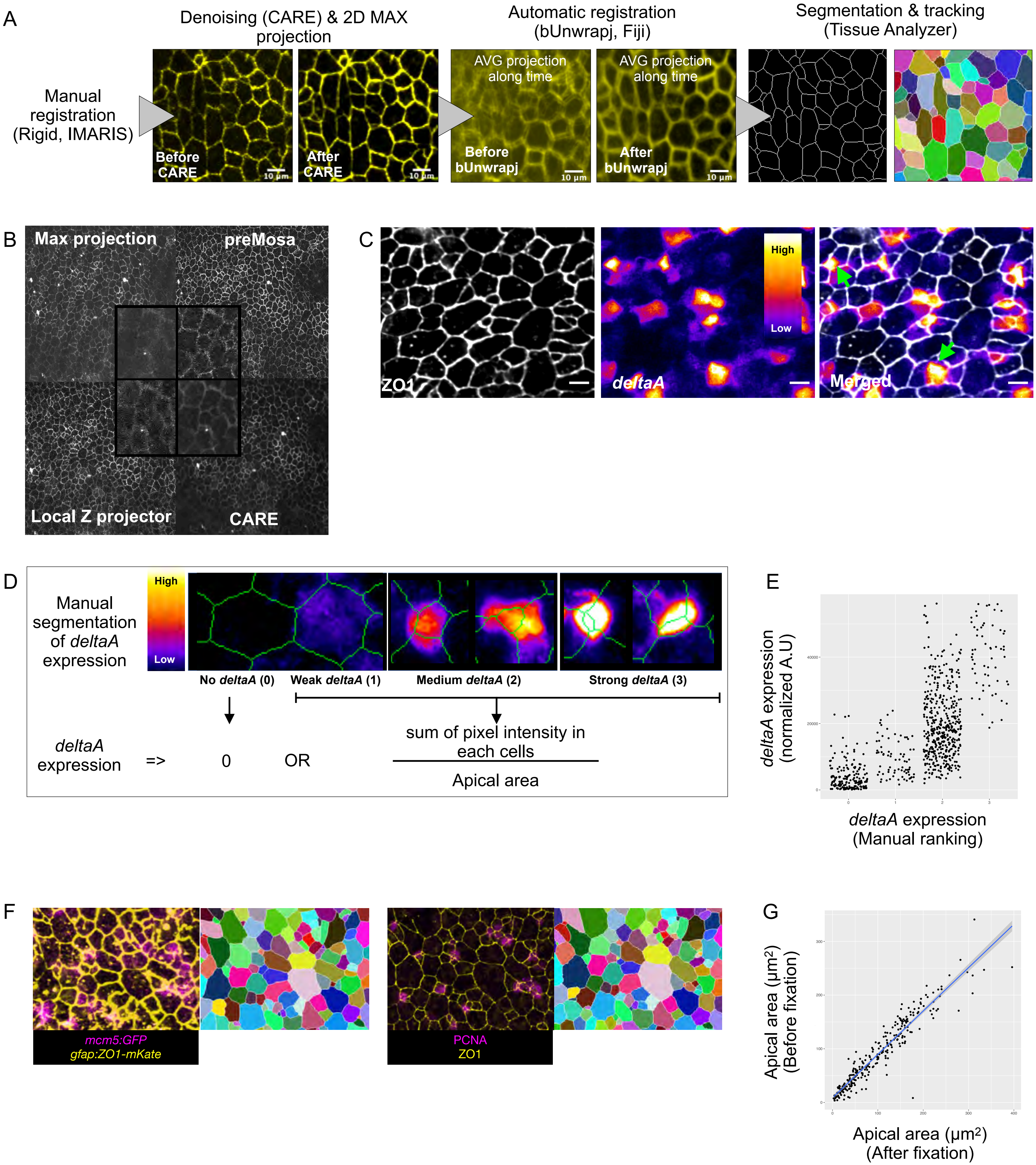

A
Manual registration (Rigid, IMARIS)
Denoising (CARE) & 2D MAX projection
Before CARE
After CARE
10 µm
Automatic registration (bUnwrapj, Fiji)
AVG projection along time
Before bUnwrapj
After bUnwrapj
Segmentation & tracking (Tissue Analyzer)
B
Max projection
preMosa
Local Z projector
CARE
C
ZO1
deltaA
High
Low
Merged
D
Manual segmentation of deltaA expression
High
Low
No deltaA (0)
Weak deltaA (1)
Medium deltaA (2)
Strong deltaA (3)
deltaA expression
=>
0
OR
sum of pixel intensity in each cells
Apical area
E
deltaA expression (normalized A.U)
deltaA expression (Manual ranking)
F
mcm5:GFP
gfap:ZO1-mKate
PCNA
ZO1
G
Apical area (µm²) (Before fixation)
Apical area (µm²) (After fixation)

### Figure S3

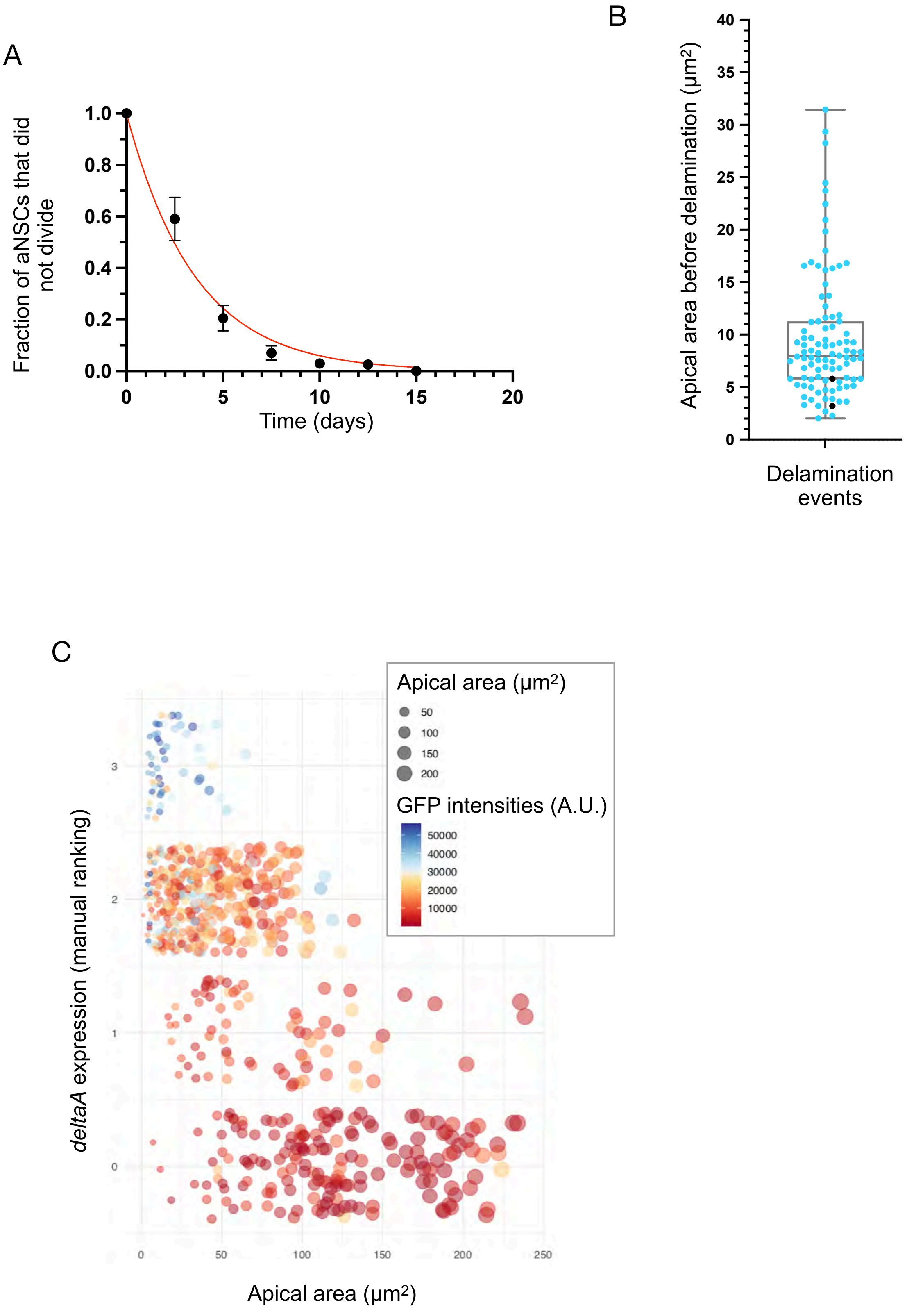
A
Fraction of aNSCs that did not divide
1.0
0.8
0.6
0.4
0.2
0.0
0
5
10
15
20
Time (days)
B
Apical area before delamination (µm2)
40
35
30
25
20
15
10
5
0
Delamination events
C
deltaA expression (manual ranking)
3
2
1
0
0
50
100
150
200
250
Apical area (µm2)
Apical area (µm2)
50
100
150
200
GFP intensities (A.U.)
50000
40000
30000
20000
10000

### Figure S3

D

*deltaA$^{neg}$* MCs

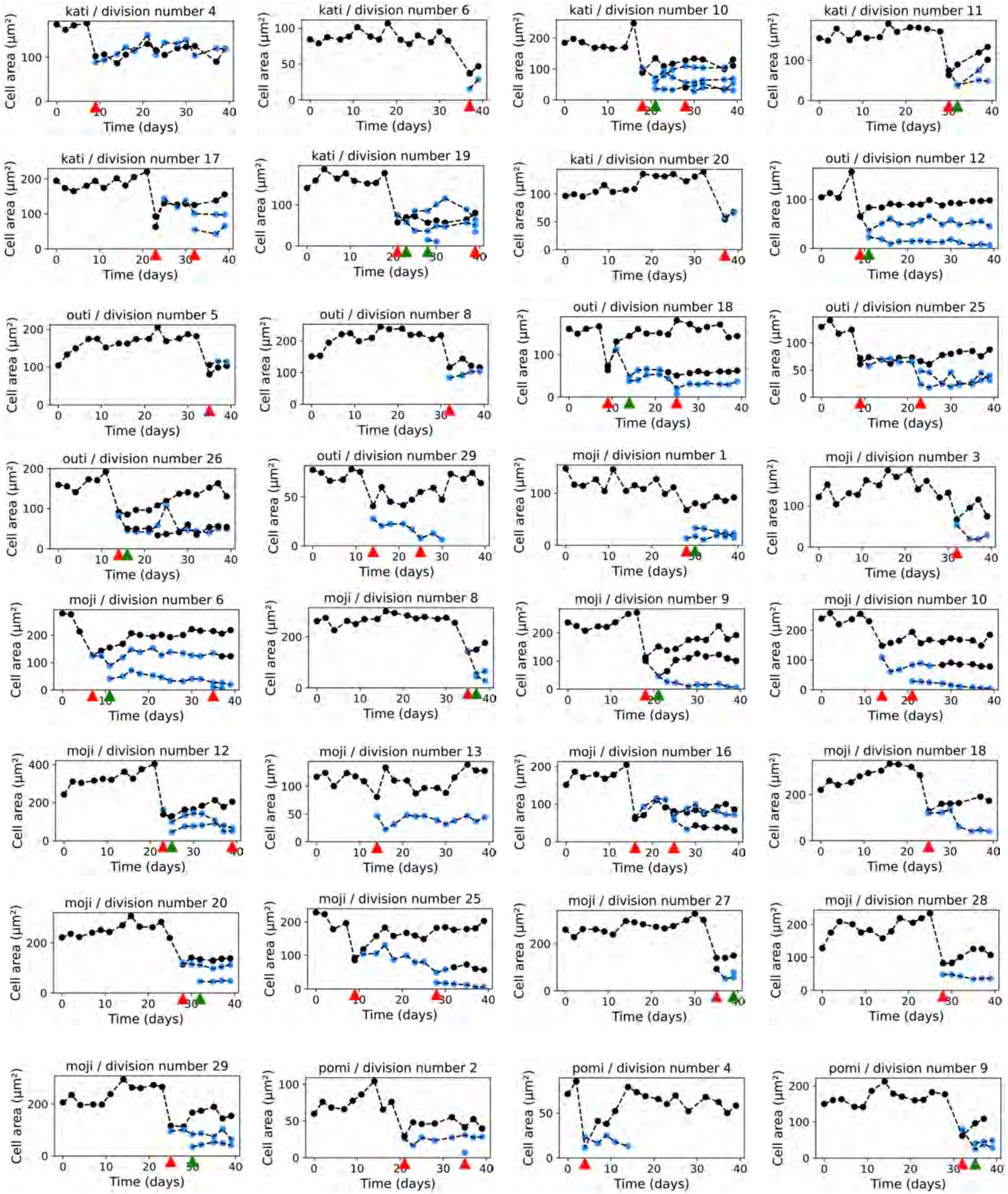

### Figure S3

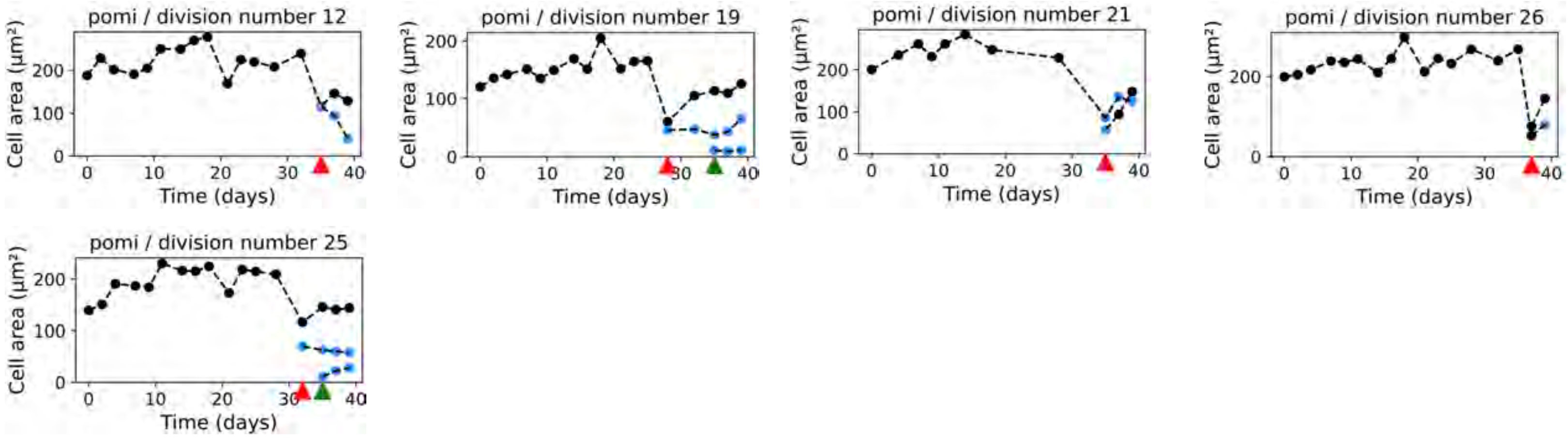


#### *deltaA*$^{pos}$ MCs

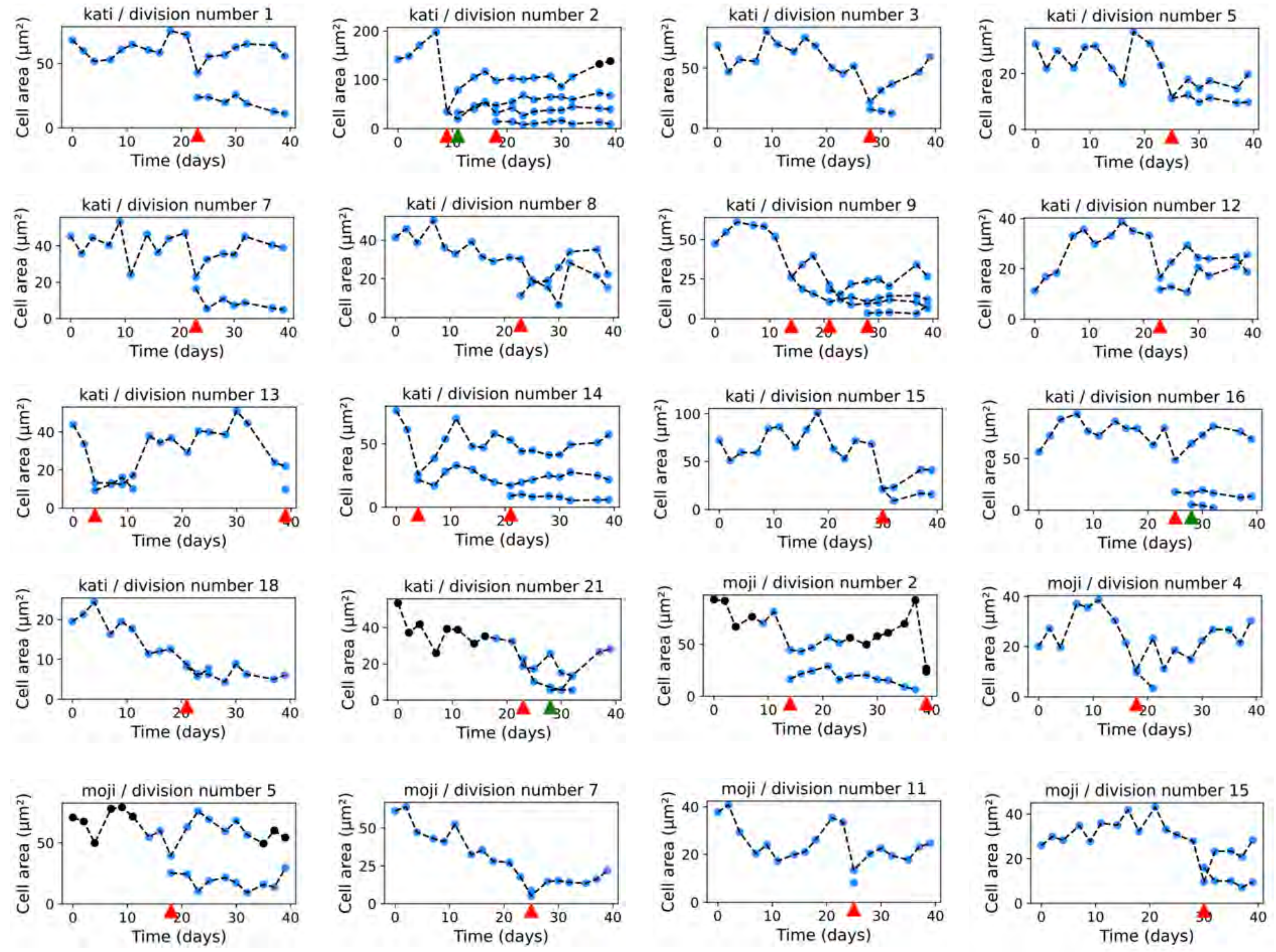

### Figure S3

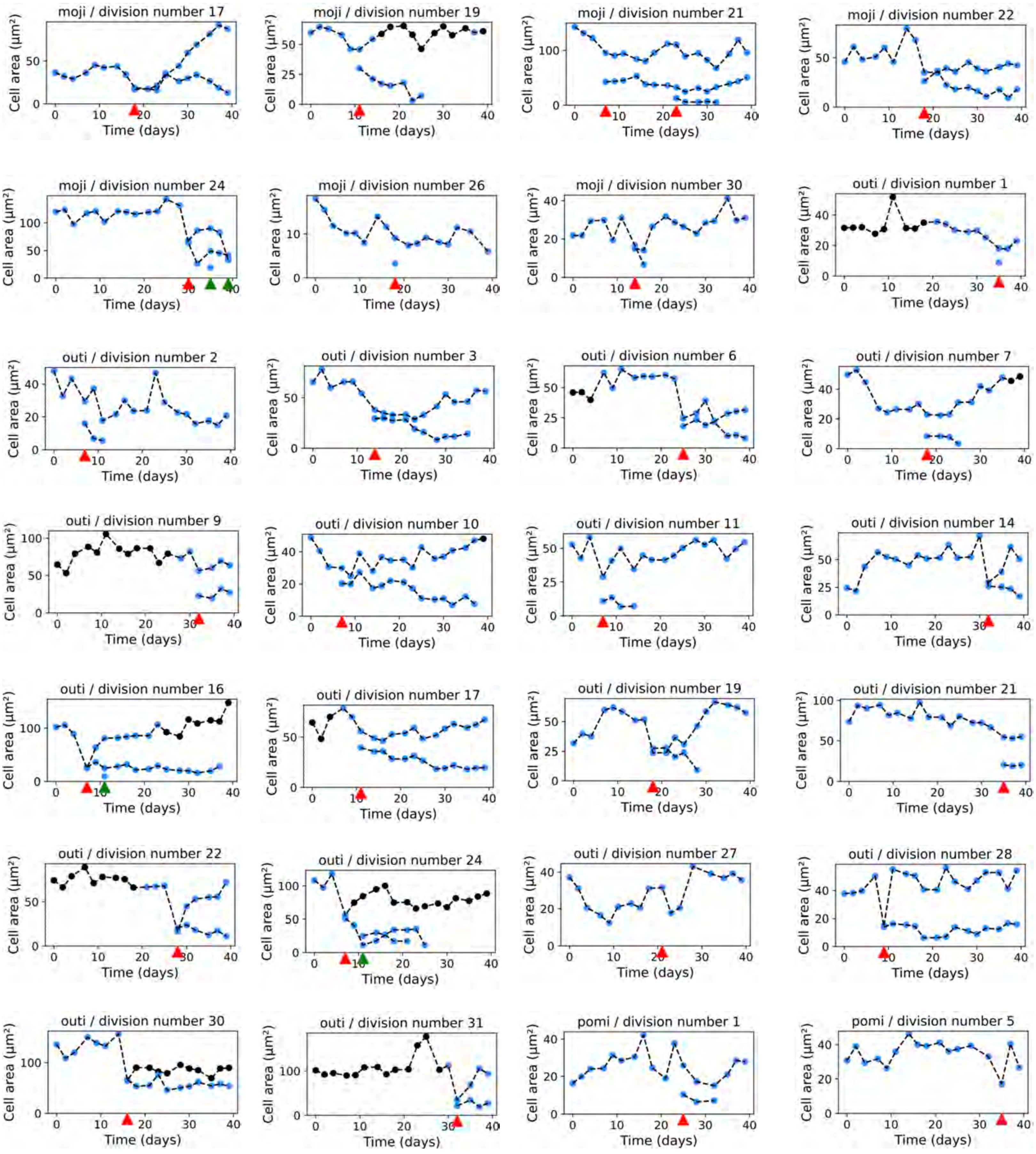
moji / division number 17
moji / division number 19
moji / division number 21
moji / division number 22
moji / division number 24
moji / division number 26
moji / division number 30
outi / division number 1
outi / division number 2
outi / division number 3
outi / division number 6
outi / division number 7
outi / division number 9
outi / division number 10
outi / division number 11
outi / division number 14
outi / division number 16
outi / division number 17
outi / division number 19
outi / division number 21
outi / division number 22
outi / division number 24
outi / division number 27
outi / division number 28
outi / division number 30
outi / division number 31
pomi / division number 1
pomi / division number 5
Cell area (µm²)
Time (days)

### Figure S3

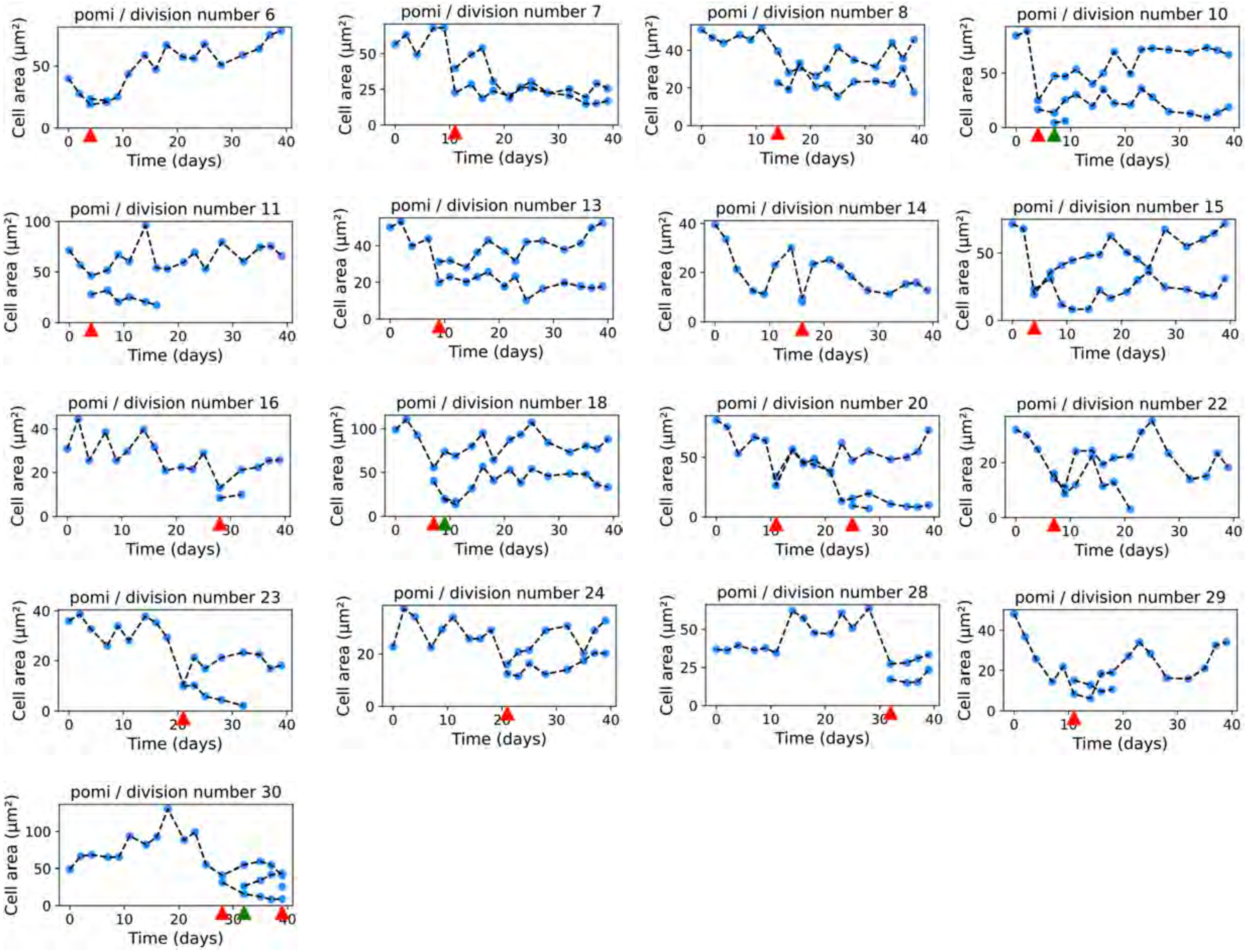
pomi / division number 6
pomi / division number 7
pomi / division number 8
pomi / division number 10
pomi / division number 11
pomi / division number 13
pomi / division number 14
pomi / division number 15
pomi / division number 16
pomi / division number 18
pomi / division number 20
pomi / division number 22
pomi / division number 23
pomi / division number 24
pomi / division number 28
pomi / division number 29
pomi / division number 30
Cell area (µm²)
Time (days)

### Figure S4

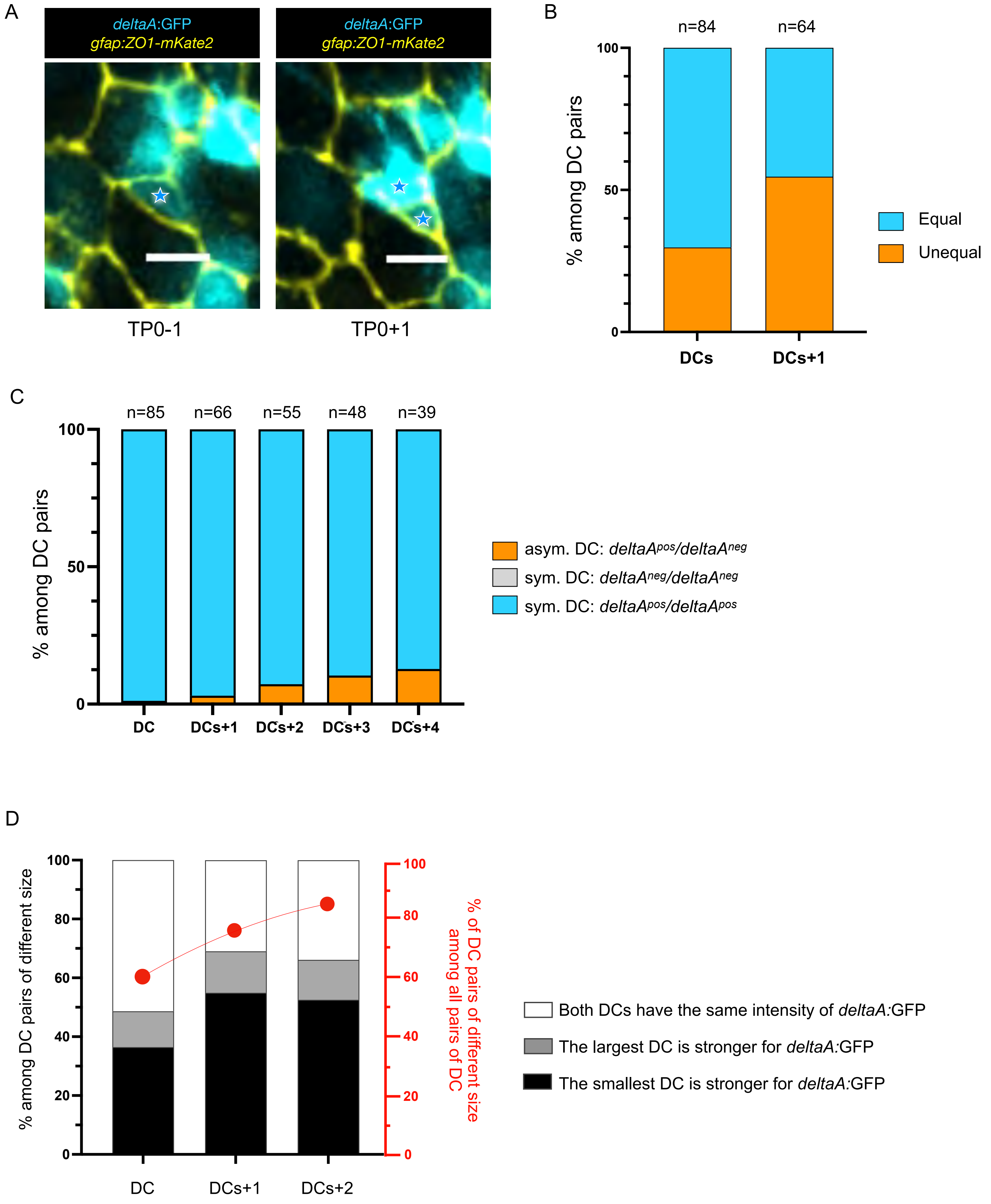

A
deltaA:GFP
gfap:ZO1-mKate2
deltaA:GFP
gfap:ZO1-mKate2
TP0-1
TP0+1
B
n=84
n=64
100
50
0
% among DC pairs
DCs
DCs+1
Equal
Unequal
C
n=85
n=66
n=55
n=48
n=39
100
50
0
% among DC pairs
DC
DCs+1
DCs+2
DCs+3
DCs+4
asym. DC: deltaApos/deltaAneg
sym. DC: deltaAneg/deltaAneg
sym. DC: deltaApos/deltaApos
D
100
80
60
40
20
0
% among DC pairs of different size
% of DC pairs of different size among all pairs of DC
DC
DCs+1
DCs+2
Both DCs have the same intensity of deltaA:GFP
The largest DC is stronger for deltaA:GFP
The smallest DC is stronger for deltaA:GFP

### Figure S5

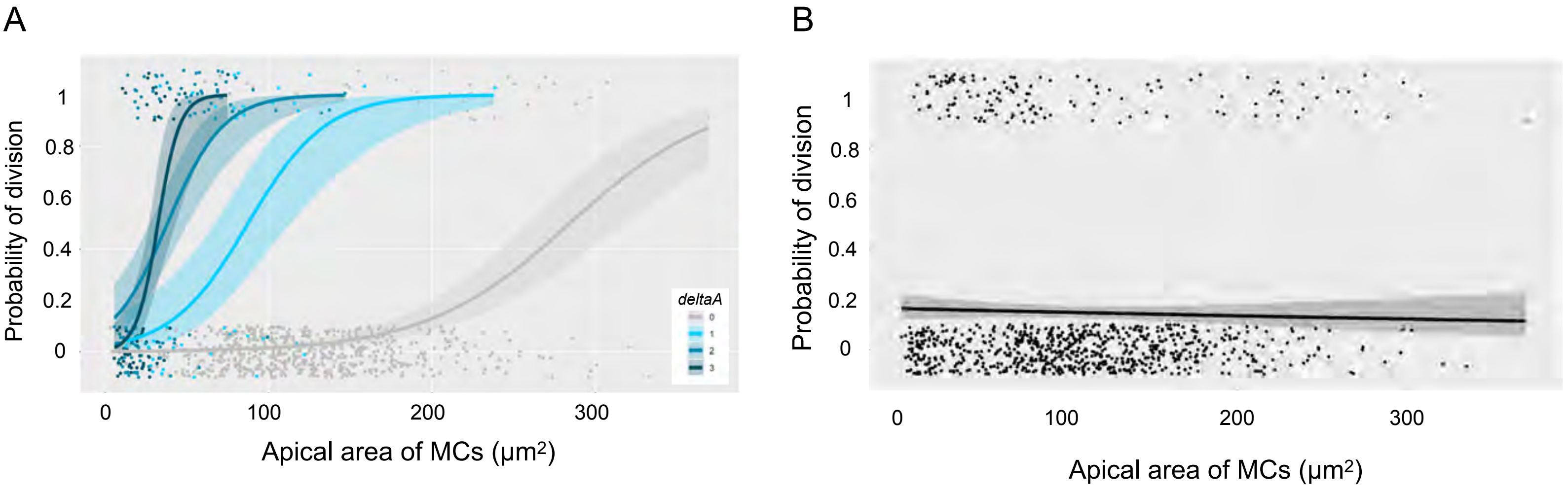
A
Probability of division
1
0.8
0.6
0.4
0.2
0
deltaA
0
1
2
3
0
100
200
300
Apical area of MCs (μm2)
B
Probability of division
1
0.8
0.6
0.4
0.2
0
0
100
200
300
Apical area of MCs (μm2)

### Figure S6

A

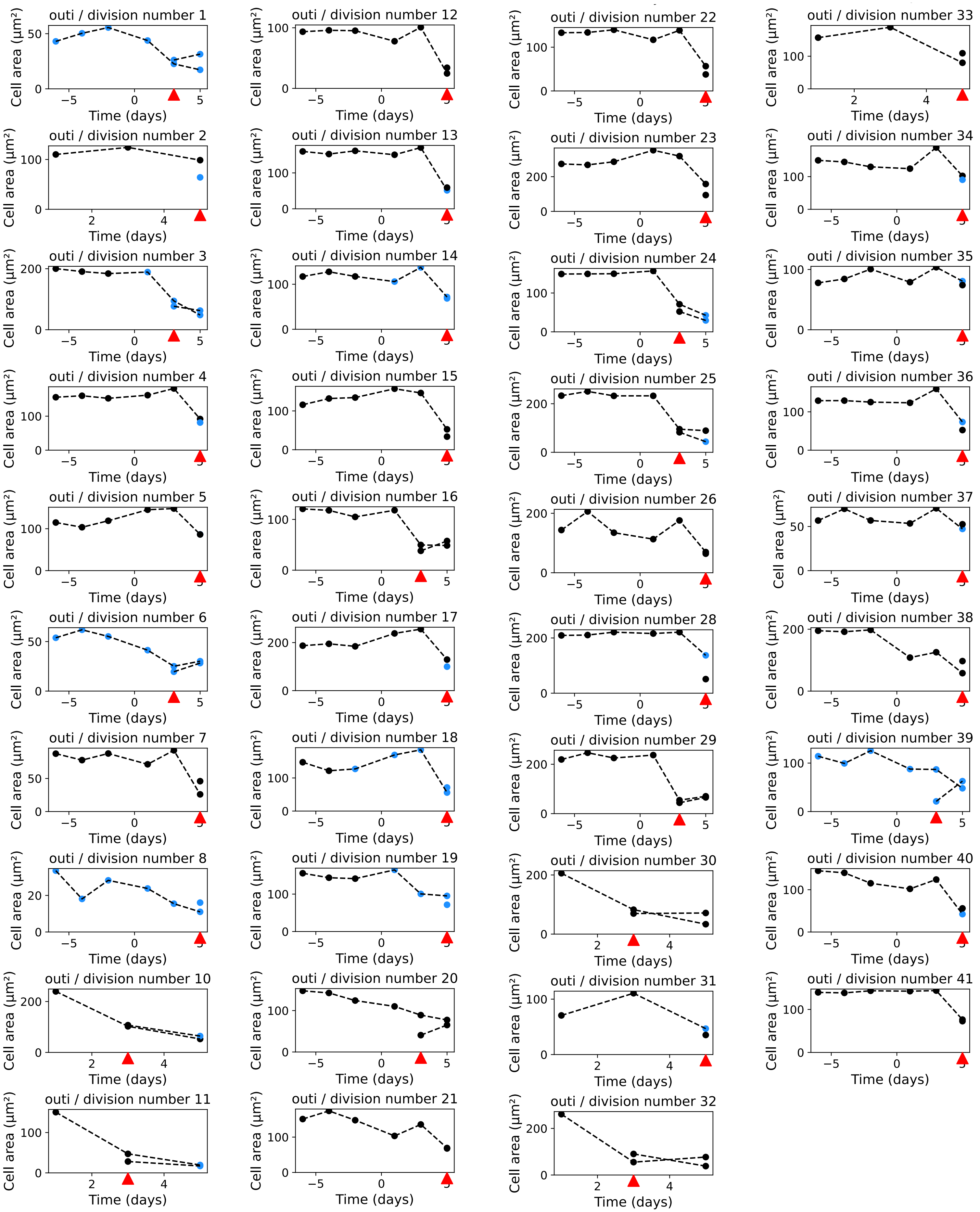
outi / division number 1
outi / division number 2
outi / division number 3
outi / division number 4
outi / division number 5
outi / division number 6
outi / division number 7
outi / division number 8
outi / division number 10
outi / division number 11
outi / division number 12
outi / division number 13
outi / division number 14
outi / division number 15
outi / division number 16
outi / division number 17
outi / division number 18
outi / division number 19
outi / division number 20
outi / division number 21
outi / division number 22
outi / division number 23
outi / division number 24
outi / division number 25
outi / division number 26
outi / division number 28
outi / division number 29
outi / division number 30
outi / division number 31
outi / division number 32
outi / division number 33
outi / division number 34
outi / division number 35
outi / division number 36
outi / division number 37
outi / division number 38
outi / division number 39
outi / division number 40
outi / division number 41
Cell area (µm²)
Time (days)

### Figure S6

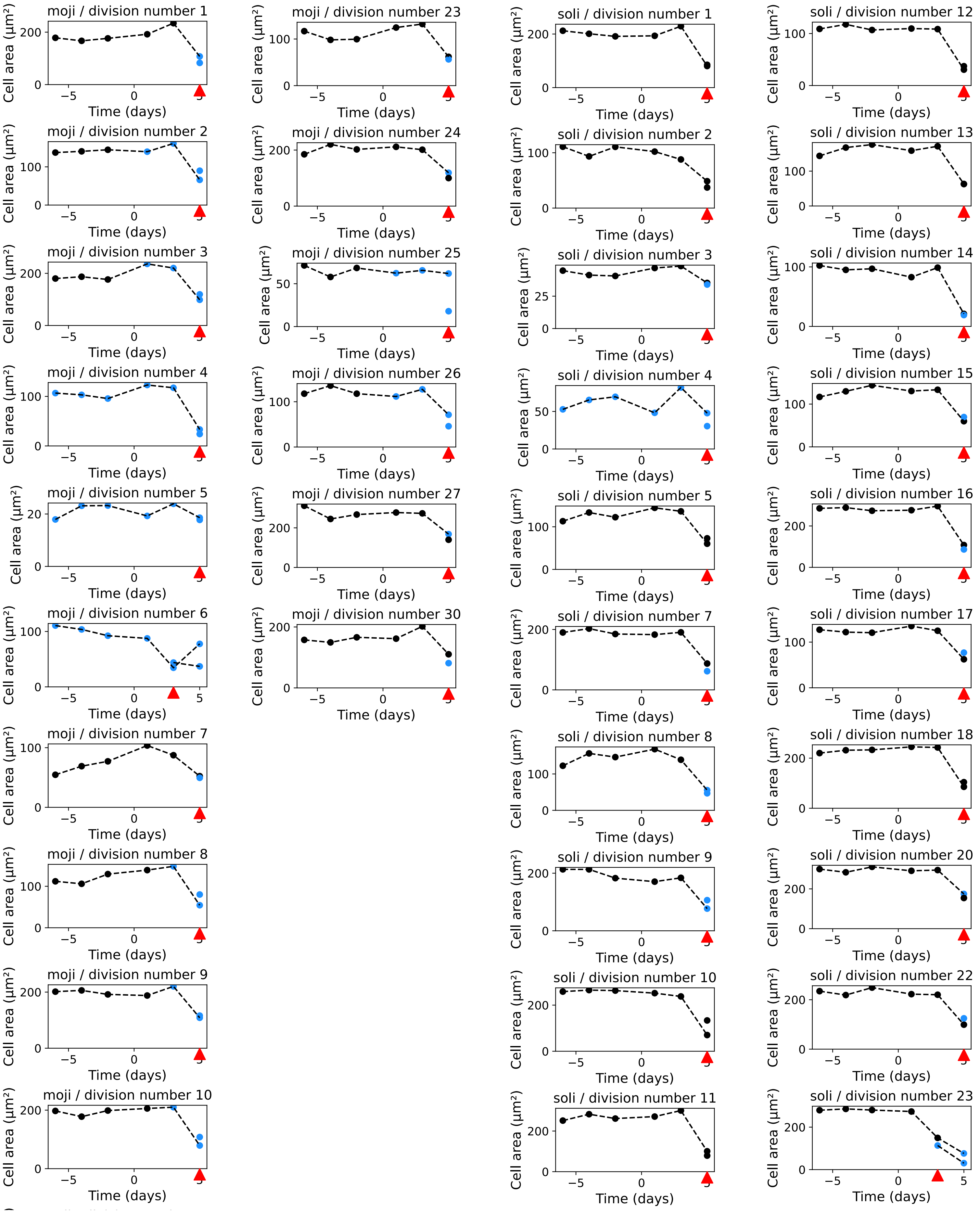
moji / division number 1
moji / division number 2
moji / division number 3
moji / division number 4
moji / division number 5
moji / division number 6
moji / division number 7
moji / division number 8
moji / division number 9
moji / division number 10
moji / division number 23
moji / division number 24
moji / division number 25
moji / division number 26
moji / division number 27
moji / division number 30
soli / division number 1
soli / division number 2
soli / division number 3
soli / division number 4
soli / division number 5
soli / division number 7
soli / division number 8
soli / division number 9
soli / division number 10
soli / division number 11
soli / division number 12
soli / division number 13
soli / division number 14
soli / division number 15
soli / division number 16
soli / division number 17
soli / division number 18
soli / division number 20
soli / division number 22
soli / division number 23
Cell area (µm²)
Time (days)

### Figure S6

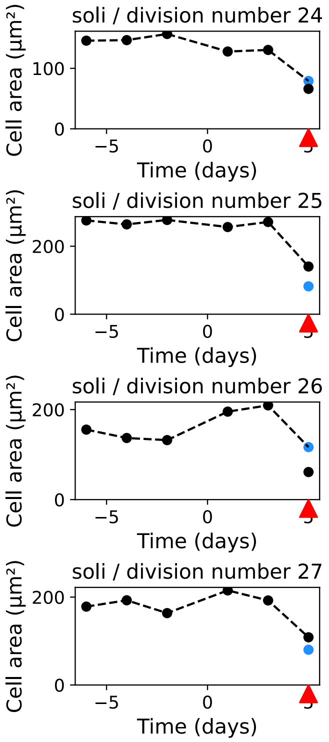
soli / division number 24
Cell area (µm²)
100
0
−5
0
5
Time (days)
soli / division number 25
Cell area (µm²)
200
0
−5
0
5
Time (days)
soli / division number 26
Cell area (µm²)
200
0
−5
0
5
Time (days)
soli / division number 27
Cell area (µm²)
200
0
−5
0
5
Time (days)

### Figure S6

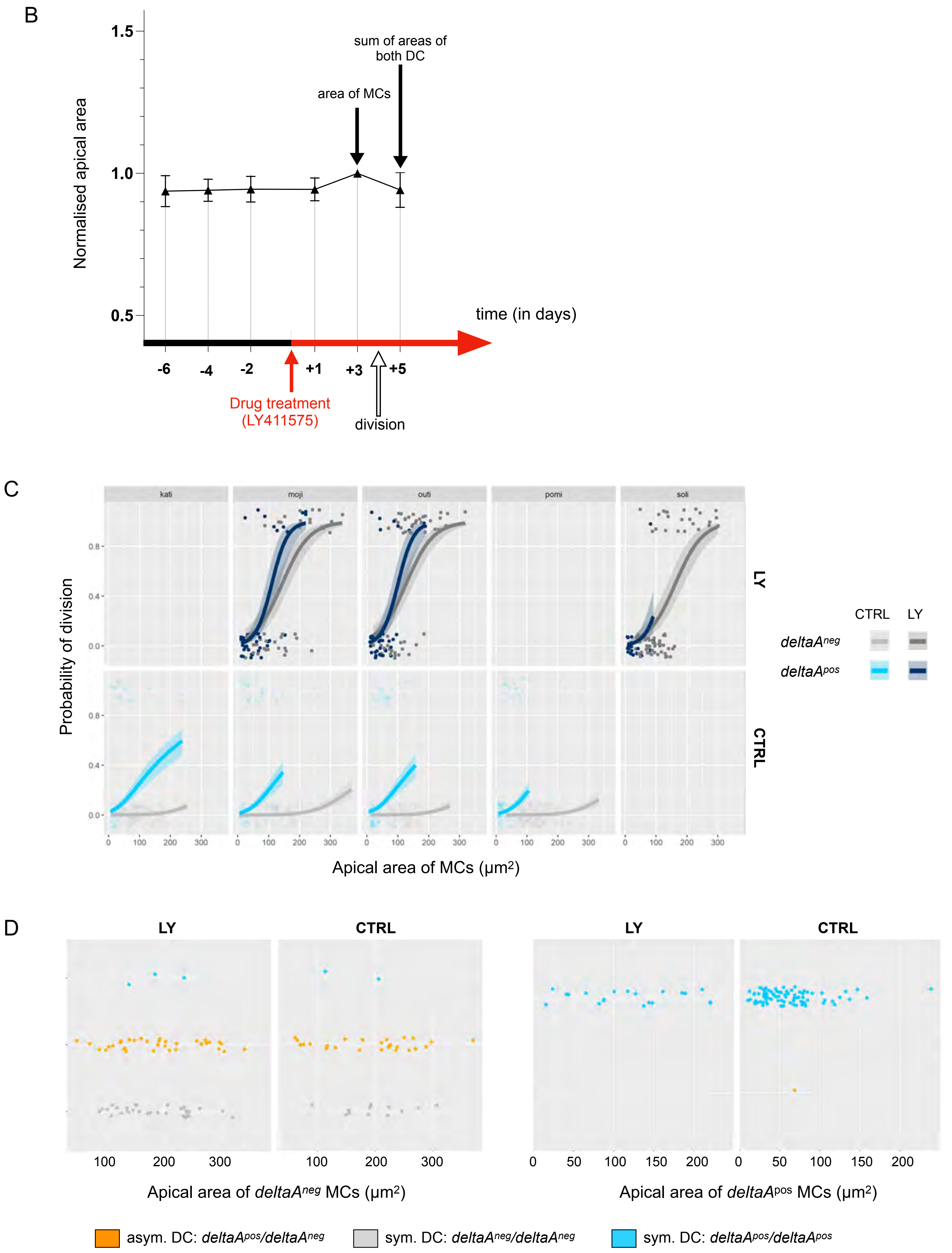
B
1.5
1.0
0.5
Normalised apical area
sum of areas of both DC
area of MCs
time (in days)
-6
-4
-2
+1
+3
+5
Drug treatment (LY411575)
division
C
kati
moji
outi
pomi
soli
LY
CTRL
Probability of division
Apical area of MCs (µm2)
CTRL
LY
deltaAneg
deltaApos
D
LY
CTRL
LY
CTRL
Apical area of deltaAneg MCs (µm2)
Apical area of deltaApos MCs (µm2)
asym. DC: deltaApos/deltaAneg
sym. DC: deltaAneg/deltaAneg
sym. DC: deltaApos/deltaApos
